# CRISPR-engineered deletion of *POGZ* alters transcription factor binding at promoters of genes involved in synaptic signaling

**DOI:** 10.1101/2025.10.27.684870

**Authors:** Mariana Moyses-Oliveira, Yating Liu, Serkan Erdin, Dadi Gao, Riya Bhavsar, Kiana Mohajeri, Kathryn O’Keefe, Philip M. Boone, Gabriela Xavier, Calwing Liao, Aiqun Li, Rachita Yadav, Monica Salani, Diane Lucente, Benjamin Currall, Celine E. F. de Esch, Derek J. C. Tai, Douglas Ruderfer, Kristen J. Brennand, James F. Gusella, Michael E. Talkowski

**Affiliations:** Center for Genomic Medicine, Massachusetts General Hospital, Boston, MA, USA; Department of Neurology, Massachusetts General Hospital and Harvard Medical School, Boston, MA, USA; Stanley Center for Psychiatric Research, Broad Institute of MIT and Harvard, Cambridge, MA, USA; Program in Medical and Population Genetics, Broad Institute of MIT and Harvard, Cambridge, MA, USA; Program in Biological and Biomedical Sciences, Harvard Medical School, Boston, MA, USA; Division of Genetics and Genomics, Boston Children’s Hospital, Boston, MA, USA; Analytic and Translational Genetics Unit, Department of Medicine, Massachusetts General Hospital, Boston, MA, USA; Department of Genetics and Genomic Sciences, Icahn School of Medicine at Mount Sinai, New York, NY, USA; Nash Family Department of Neuroscience, Icahn School of Medicine at Mount Sinai, New York, NY, USA; Mount Sinai Center for Transformative Disease Modeling, Icahn School of Medicine at Mount Sinai, New York, NY, USA; Friedman Brain Institute, Icahn School of Medicine at Mount Sinai, New York, NY, USA; Icahn Institute for Data Science and Genomic Technology, Icahn School of Medicine at Mount Sinai, New York, NY, USA; Division of Genetic Medicine, Department of Medicine, Vanderbilt Genetics Institute, Vanderbilt University Medical Center, 1211 Medical Center Dr. Nashville, TN, USA; Department of Biomedical Informatics and Department of Psychiatry and Behavioral Sciences, Vanderbilt University Medical Center, 1211 Medical Center Dr. Nashville, TN, USA; Department of Psychiatry, Yale University New Haven, CT, USA; Department of Genetics, Blavatnik Institute, Harvard Medical School, Boston, MA, USA; Harvard Stem Cell Institute, Harvard University, Cambridge, MA, USA

**Author notes:** These authors contributed equally (Mariana Moyses-Oliveira, Yating Liu, Serkan Erdin, Dadi Gao).

## Abstract

One of the seminal discoveries from genetic studies of autism spectrum disorder and related neurodevelopmental disorders (NDD) has been that loss-of-function (LoF) mutations in many genes that impact chromatin and transcriptional regulation confer substantial liability to NDD. Haploinsufficiency of the epigenetic regulator *POGZ* represents one of the strongest such associations; however, little is known about the direct or indirect regulatory targets of *POGZ*, or the mechanisms by which loss of this chromatin modifier alters early neuronal development and synaptic functions. Here, we created an allelic series of CRISPR-engineered human induced pluripotent stem cell (hiPSC) clones harboring mono- and biallelic *POGZ* deletions. In hiPSC-derived neural stem cells (NSC) and Neurogenin 2-induced neurons (iN), *POGZ* LoF altered the expression of genes associated with critical cellular processes and neuronal functions, including synaptic and intracellular signaling and extracellular matrix organization. Our multiomics profiling also showed altered footprinting of critical transcription factors (e.g., activator protein 1 complexes) that were enriched at promoters of differentially expressed genes associated with synaptic function. To further interrogate the shared molecular changes in neuronal development associated with NDD and *POGZ* regulation, we compared our results to deletions of the transcription factor MEF2C and the sodium channel gene SCN2A that we generated in these same isogenic iN. These analyses revealed strong enrichment of extracellular matrix and intracellular signaling disruption associated with *POGZ* and MEF2C deletion, whereas *POGZ* and SCN2A haploinsufficiency exhibited shared transcriptional effects on gene modules enriched for NDD-associated genes with opposing regulatory effects. Notably, we also observed alterations to synaptic firing rate and neurite extension with biallelic deletions, but not heterozygous lines, suggesting subtle effects in neuronal development associated with haploinsufficiency. Overall, these shared molecular consequences suggest key points of convergence that connect epigenetic regulation to neuronal function in the etiology of neurodevelopmental pathologies.

## INTRODUCTION

Remarkable progress in recent years has identified shared genetic etiologies across a broad spectrum of neurodevelopmental disorders (NDD), including autism spectrum disorder (ASD), developmental delay, intellectual disability, and many diverse developmental syndromes^1–4^. Common hallmarks of these gene discovery efforts have been the consistently reproducible association of *de novo* loss-of-function (LoF) mutations in genes highly intolerant to functional variation in the general population (i.e., constrained) and directly involved in pathways governing synaptic activity and gene regulation (e.g., transcriptional regulation and chromatin organization)^1,5–9^. The primary challenge is now to connect the mechanisms by which alterations to genes involved in broad genome regulatory functions such as chromatin modification are interconnected, either directly or indirectly, with those genes involved in synaptic activity and collectively result in comparable deficits in neuronal function^9–11^. Some LoF mutations in chromatin-related genes can influence synaptic activity by dysregulating neuronal gene expression, or through other mechanisms such as proximal interactions, post-translational modifications, protein-protein interactions, or signaling pathways^7,9,11–17^. However, the broader convergence of chromatin and synaptic mechanisms, as well as their involvement in neurodevelopment and NDD pathogenesis remain uncertain.

*POGZ* (Pogo transposable element derived with ZNF domain) is among the chromatin-related genes with the most robust statistical evidence for association with ASD and more broadly defined NDD^1,7^. Individuals with heterozygous LoF mutations in *POGZ* often present with ASD, intellectual disability (ID), and/or White-Sutton syndrome^18,19^. Mouse models with homozygous *Pogz* LoF or expression of a *POGZ* patient-specific missense mutation show impairments in neurogenesis, neurodifferentiation, neuronal migration, and electrophysiological activity^20,21^. However, little is known concerning POGZ’s role in human neurodevelopment. Biochemical studies have shown that POGZ forms complexes with other ASD/NDD-associated chromatin regulators, e.g., ADNP, CHD4, and the BAF complex^22,23^. Also, the POGZ-interacting protein, heterochromatin protein 1 (HP1), is known to modulate chromatin topology in pericentromeric heterochromatin regions and euchromatic sites^24,25^. Although POGZ was first described as a transcriptional repressor through its interaction with HP1^26,27^, it has been suggested recently that POGZ might modulate gene expression as both a repressor and an activator^21,22^. Consequently, the impact of *POGZ* disruption on neuronal function and potential convergence with the disruption of genes involved in synaptic activity are fundamental insights that are required to gain understand the POGZ-associated neurodevelopmental syndromes and traits.

We took a systematic neuronal modeling approach to interrogate the impact of *POGZ* haploinsufficiency on cellular functions in developing neural cells. We engineered an allelic series of isogenic human induced pluripotent stem cell (hiPSC)-derived neural stem cell (NSC) and Neurogenin 2 (*Ngn2*)-induced neuron (iN) models that harbored CRISPR generated deletions of POGZ. We used heterozygous lines to explore the transcriptional, regulatory, and cellular consequences of *POGZ* haploinsufficiency. Additional lines with homozygous *POGZ* knockout grounded validation and mechanistic analysis of POGZ function. We then compared transcriptional signatures of *POGZ* LoF to the effects of deletions engineered on the same isogenic backgrounds for two well-established ASD/NDD genes associated with *POGZ* regulatory changes — *MEF2C*, which encodes a transcriptional regulator, and *SCN2A*, a sodium channel gene. Our analyses disclosed functional signatures implicating direct and indirect *POGZ* regulatory changes that altered transcription factor binding at the promoters of synaptic proteins and shared molecular consequences with *MEF2C* and *SCN2A* that converged at the level of genes associated with altered neuronal development.

## RESULTS

### Generation of an allelic series of isogenic neural cell lines harboring POGZ deletions

We created *POGZ* CRISPR-mediated LoF models on two different hiPSC backgrounds (GM08830 [male] and MGH2069 [female]), including a full and a partial deletion of *POGZ*, and corresponding unedited wild-type (WT) models that were exposed to the same CRISPR guide RNAs and single-cell sorting (**Figure 1A, Supplementary Figure 1A and 1B**). We differentiated these lines into two distinct neural cell types relevant to neuronal development and function, i.e., NSC and iN (**Figure 1B**), and evaluated transcriptional profiles by RNA sequencing (RNA-seq) and chromatin accessibility by the assay for transposase-accessible chromatin using sequencing (ATAC-seq) (**Figure 1C**). We generated a total of 66 RNA-seq libraries plus 66 ATAC-seq libraries for each cell type (NSC and iN), derived from 66 hiPSC differentiation replicates. The differentiation replicates were originated from a total of 23 and 24 independently CRISPR-engineered clonal hiPSC lines clones in GM08830 and MGH2069 backgrounds, respectively. We performed differential expression analyses after validating *POGZ* dosage and downregulation of RNA and protein expression in the edited lines (**Supplementary Figure 1C-F**). Comparison of the heterozygous *POGZ* LoF lines to the matched unedited isogenic control lines discovered differentially expressed genes (DEG) resulting from *POGZ* haploinsufficiency. In contrast, lines with complete loss of POGZ due to compound heterozygous LoF mutations provided validation and further mechanistic insights. In all experiments, we assessed DEG for biological significance using a false discovery rate (FDR) q<0.1 threshold and concordant direction of effect across two different hiPSC backgrounds (e.g., consistently upregulated or downregulated DEG in GM08830 and MGH2069 backgrounds; see **Supplemental information** for details).

**Figure 1.**
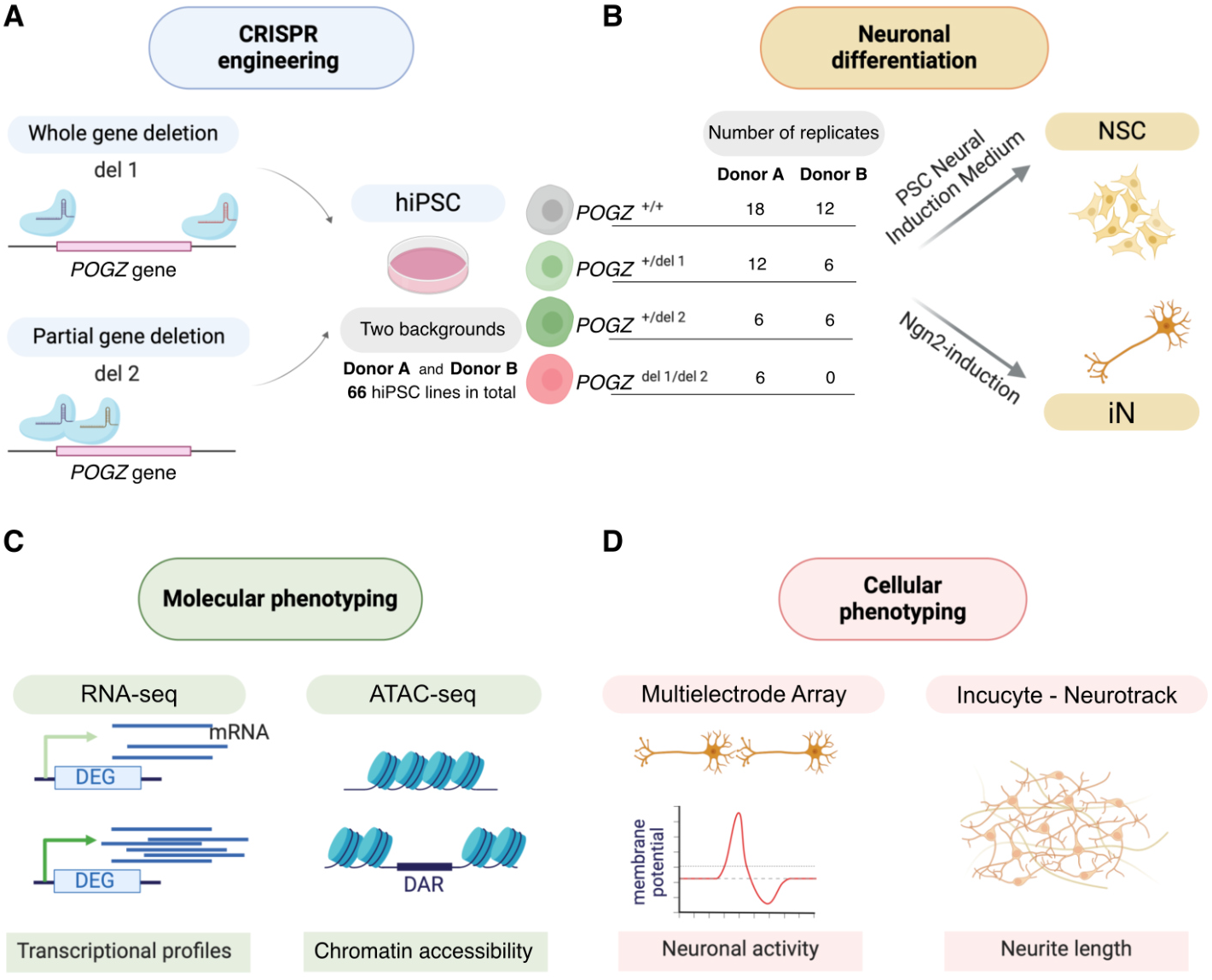
Characterization of the consequences of POGZ loss of function in neuronal models. **(A)** CRISPR-engineering of whole gene deletion (del 1) and partial gene deletion (del 2) of POGZ gene in two different backgrounds of human induced pluripotent stem cells (hiPSC) (GM08830 [Donor A, male] and MGH2069 [Donor B, female]), resulting in heterozygous (het) and compound heterozygous (comp het) clones. **(B)** Neuronal differentiation of hiPSC clones into neural stem cells (NSC) and Ngn2-induced neurons (iN). **(C)** Molecular profiling of hiPSC-derived iN and NSC using RNA-seq to identify differentially expressed genes (DEG) and ATAC-seq to address differentially accessible regions (DARs). **(D)** Cellular phenotyping of iN with multielectrode array to measure firing activity and incucyte to measure neurite length.

### POGZ haploinsufficiency is associated with altered expression of genes related to extracellular matrix and synaptic processes

We first focused on heterozygous *POGZ* deletions to gain insight into NDD-associated *POGZ* haploinsufficiency. Neuronal lines differentiated from the different hiPSC backgrounds showed significant directionally-concordant overlap of DEG across the backgrounds (GM08830 and MGH2069) in the transcriptional effects of heterozygous *POGZ* deletion (**Supplementary Figure 2A-D**). In heterozygous iN, we observed 424 DEG (146 upregulated and 278 downregulated; **Figure 2A, Supplementary Table 1**) that were significantly enriched for molecular pathways related to synapse activity and extracellular matrix, calcium signaling, and cell migration, among others (**Figure 2B**). In heterozygous NSC, the 1,115 DEG (664 upregulated and 451 downregulated; **Figure 2A, Supplementary Table 2**) were significantly enriched for similar extracellular matrix and focal adhesion-related pathways (**Figure 2B**). There was also robust statistical evidence for overlap between the DEG in iN and NSC (Fisher’s exact test, p-value=1.14e-10; **Supplementary Figure 2E and F**). Most significantly enriched pathways among iN and NSC DEG comprised a mixture of downregulated and upregulated genes (**Supplementary Figure 3A**), suggesting that the functional consequences of *POGZ* perturbation involve both transcriptional activation and repression.

**Figure 2.**
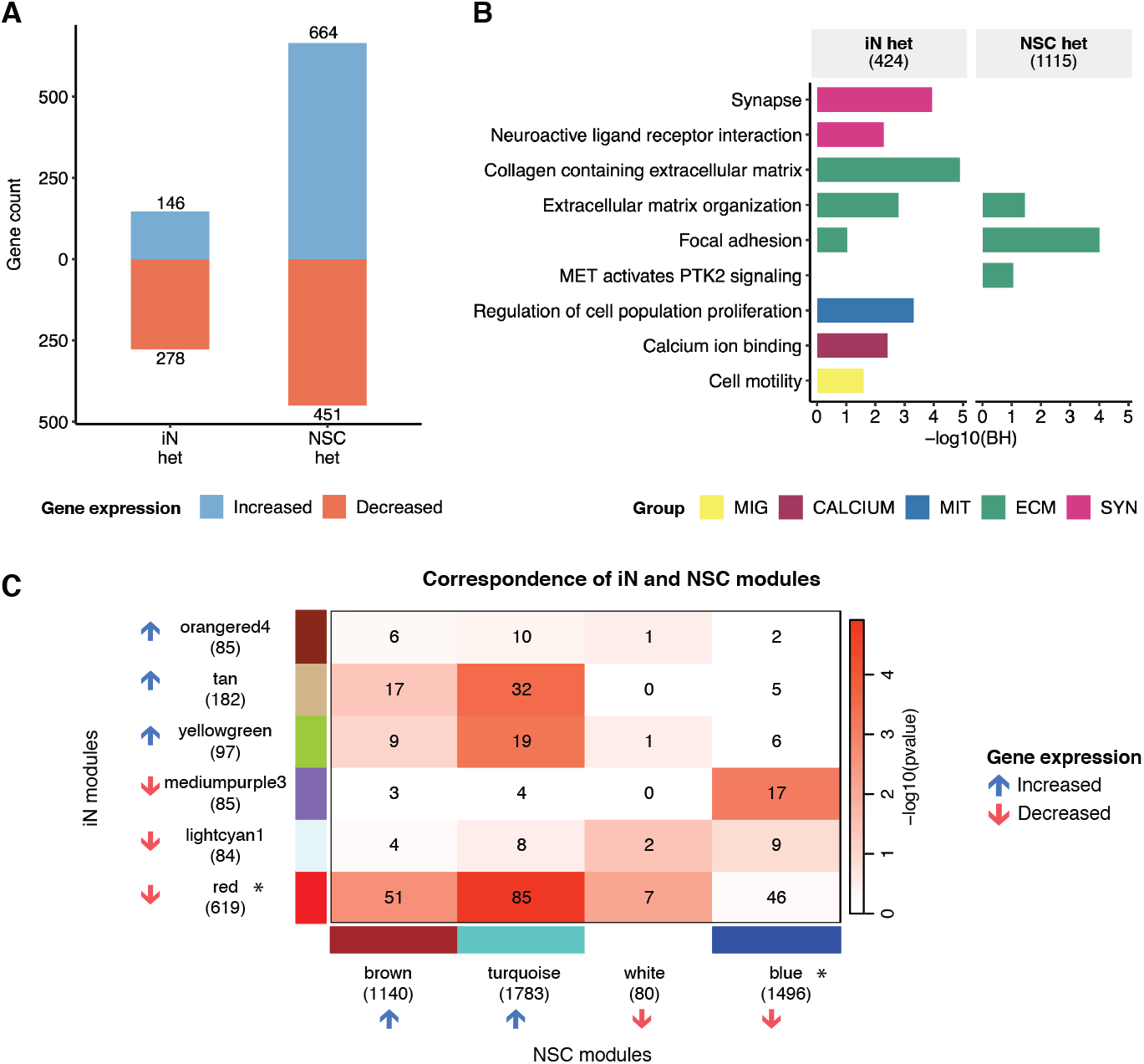
Transcriptional profiles of NSC and iN lines with heterozygous POGZ deletion. **(A)** Number of upregulated (blue) and downregulated (red) genes in iN and NSC heterozygous (het) models. **(B)** Statistical significance for enrichment of pathways among DEG in heterozygous (het) N and NSC lines. Numbers in parentheses indicate the total number of genes in a given gene list. Colors represent pathway categorization according to the nature of the functional categories related to the enriched term. Groups of functional categories: VESIC: vesicles, ENERGY: energy metabolism, MENTAL: mental and behavioral traits, IMMUNE: immune system and inflammation, MIG: cell migration, CALCIUM: calcium signaling, MIT: mitosis and cell proliferation, STEROID: steroid metabolism, CSKEL: cytoskeleton, CHROM: chromatin structure and regulation, ECM: extracellular matrix, SYN: synapse. C. Heatmap indicating significant overlap between co-expression modules in iN and NSC. Asterisks indicate the modules containing POGZ.

Next, we established modules of co-expressed genes in NSC and iN by using weighted gene co-expression network analysis (WGCNA)^28^. For each cell type, we merged the data across both backgrounds and all deletions models (see Supplemental information for details). In WGCNA of heterozygous and control iN, six co-expression modules were significantly correlated with *POGZ* genotype (Bonferroni-adjusted p-value<0.1); three displayed downregulation and three upregulation of the module eigengene (**Supplementary Figure 3B**). In the heterozygous and control NSC, four co-expression modules were significantly correlated with *POGZ* genotype (Bonferroni-adjusted p-value<0.1), with two having eigengenes indicative of downregulation and upregulation, respectively (**Supplementary Figure 3C**). All DEG and genotype-correlated co-expression modules in both NSC and iN were significantly enriched for DEG associated with *Pogz* dosage in previously published mouse models (**Supplementary Figure 4A and B**), indicating reproducibility of *POGZ* LoF impacts across species.

*POGZ* was a member of the downregulated modules “red” (619 genes) and “blue” (1,496 genes) in iN and NSC, respectively (**Supplementary Figure 3B and C, Supplementary Table 3**). The pathway enrichment analysis of these two modules showed a strong involvement of *POGZ* co-expressed genes in cell type-specific functional categories. The iN red module was significantly enriched for processes related to extracellular matrix and growth factor binding (FDR=1.50e-6, FDR=4.43e-2, respectively), and the NSC blue module was significantly enriched for pathways associated with translation, ribosomal RNA processing and nervous system development (FDR=1.83e-168, FDR=2.18e-61, FDR=2.52e-30, respectively) (**Supplementary Figure 3D**). These modules shared 46 genes, an overlap that did not reach significance (p-value=1.93e-1, **Figure 2C**); however, the *POGZ-* containing downregulated iN red module showed significant overlap with NSC upregulated modules, “turquoise” (overlap of 85 genes, p-value=1.22e-5) and “brown” (overlap of 51 genes, p-value=2.03e-3) (**Supplementary Table 4, Figure 2C**). These gene overlaps were significantly enriched for enzyme-receptor signaling, growth factor binding, extracellular matrix organization and neurodevelopmental pathways that were also captured in the *POGZ*-containing modules (**Supplementary Figure 3D**). Thus, these transcriptional profiles indicate that *POGZ* LoF results in gene expression disturbances that can be cell-type specific or shared but directionally discordant yet converge on common cellular processes.

### POGZ direct regulatory targets modulated in edited NSC are associated with chromatin regulation, cell cycle, and RNA processing

Gene expression differences in our neuronal models could be due to direct transcriptional regulation of DEG by POGZ binding at the promoter (see definitions of POGZ direct effects in **Supplementary Figure 5A**). To interrogate the modulation of genes that are direct targets of POGZ, we compared the iN and NSC DEG with POGZ targets from published CUT&RUN or CUT&Tag experiments defining POGZ binding sites in human fetal cortex ^22^ (human POGZ targets) and mouse embryonic stem cells (mESC)^23^ (mouse Pogz targets), respectively (**Supplementary Table 5**).

Downregulated DEG from heterozygous NSC lines overlapped significantly with the human POGZ targets (p-value=1.06e-4), and both the upregulated and downregulated NSC DEG showed mild enrichment of mouse Pogz targets (**Supplementary Figure 6A**). The smaller number of DEG from heterozygous iN lines showed no such enrichments (**Supplementary Figure 6A)**. Across the WGCNA co-expression modules correlated with *POGZ* genotype, only the downregulated NSC blue module containing *POGZ* showed strong enrichment of POGZ target genes (**Supplementary Figure 6B**). These 498 overlapping genes were significantly enriched for DNA binding, transcription, and RNA processing-associated processes (**Supplementary Figure 6C**). The same pathways were also detected by the 145 genes shared between NSC downregulated DEG and human POGZ targets (**Supplementary Figure 6C**), arguing that one disrupted function in NSC harboring POGZ haploinsufficiency was the role of activating expression of some direct target genes involved in these biological processes. However, these analyses also argue more broadly that most gene expression changes caused by *POGZ* heterozygous disruption do not result from direct regulation by POGZ binding at the promoter of the DEG, but rather indirect regulatory changes.

### Complete inactivation of POGZ impacts activity of its direct target genes

We next sought to confirm these new insights into *POGZ* function by investigating cell lines with complete inactivation due to compound heterozygous mutations. The strong transcriptional consequences (9,006 and 5,148 DEG in iN and NSC, respectively, **Supplementary Figure 7A, Supplementary Table 6 and 7**) were highly correlated with the impacts of heterozygous LoF mutations (**Supplementary Figure 7B-F**). Similarly to our findings for iN and NSC co-expression modules, subsets of DEG overlapped between different zygosities and cell types (**Supplementary Figure 7B and C**). Some of these DEG overlaps presented with discordant directions (e.g., downregulated in heterozygous iN and upregulated in compound heterozygous NSC) (**Supplementary Figure 7B**).

In compound heterozygous iN, POGZ direct targets (**Supplementary Figure 7G**) overlapped significantly with upregulated DEG (p-value=1.48e-23), implicating POGZ as a transcriptional repressor in this cell type. These POGZ direct target DEG overlapped significantly with NDD genes previously implicated by large-scale exome studies (**Supplementary Figure 7H**), but this pattern was not observed in the context of heterozygous mutations (**Supplementary Figure 7I**). The NDD-associated direct POGZ target genes upregulated in iN showed significant enrichment for pathways related to synapse, immune, cell cycle, and translation regulation processes (**Figure 3**). Similar functional categories were recapitulated by the pathways enriched among NDD-associated direct POGZ target genes downregulated in NSC (**Figure 3**). These analyses indicate that *POGZ* directly modulates the transcription of other NDD-associated genes.

**Figure 3.**
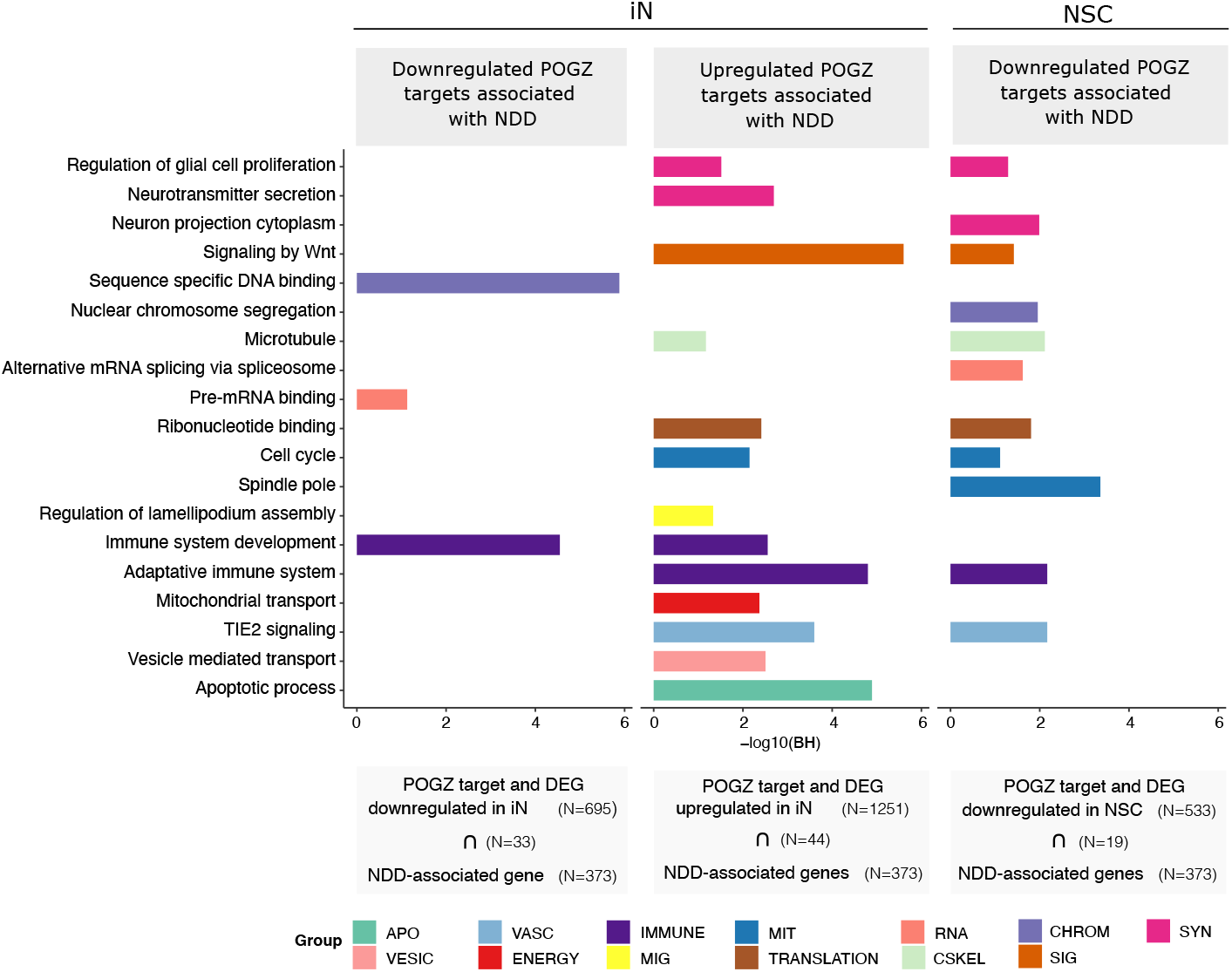
Impacts of compound heterozygous POGZ deletion on ASD and NDD-associated genes. Statistical significance for enrichment of pathways among NDD-associated genes that are downregulated (left panel) or upregulated (middle panel) differentially expressed POGZ targets in iN compound heterozygous lines. Statistical significance for enrichment of pathways among NDD-associated genes that are downregulated differentially expressed POGZ targets in NSC compound heterozygous lines (right panel). Intersection of two gene lists is represented by “⋂”. Numbers in parentheses indicate the total number of genes in a given gene list. Colors represent pathway categorization according to the nature of the functional categories related to the enriched term. Groups of functional categories: APO: apoptosis, VESIC: vesicles, VASC: vascular system, ENERGY: energy metabolism, IMMUNE: immune system and inflammation, MIG: cell migration, MIT: mitosis and cell proliferation, TRANSLATION: translation, RNA: RNA metabolism, STEROID: steroid metabolism, CSKEL: cytoskeleton, CHROM: chromatin structure and regulation, SIG: intracellular signaling pathways, ECM: extracellular matrix, SYN: synapse. BH: Benjamini-Hochberg adjusted p-values.

### POGZ’s interactors influence the direction of its transcriptional modulation

One potential explanation for cell type-dependency of directional differences in the regulation of POGZ targets pertains to the protein(s) with which POGZ is complexed when it binds to DNA. POGZ is known to form a variety of complexes with chromatin proteins, such as HP1, as well as other NDD risk-associated transcription regulators, such as ADNP, CHD4, and BRG1, the latter being a member of the BAF complex, which has also been implicated in NDD^21–23,29^. To investigate the influence of POGZ’s interactors on expression modulation, we compared NSC and iN DEG with lists of mouse Pogz targets defined with respect to the composition of the POGZ-containing protein complex (See methods for details)^23^. Importantly, all such subsets of mouse Pogz targets showed significant overlap with POGZ targets in the human fetal cortex (**Supplementary Figure 8A**), and the mono- and biallelically edited lines demonstrated concordant DEG directionalities in the context of POGZ complex composition (**Supplementary Figure 9**).

As expected, compound heterozygous iN lines revealed a strong correlation between upregulated DEG and genomic targets of multiple POGZ complex configurations (**Supplementary Figure 8B**), reinforcing POGZ’s important role as a repressor of gene expression in these cells, though a significant subset of Pogz/Chd4 targets were downregulated, consistent with an activator role. In NSC compound heterozygous lines, genes matching mouse Pogz/ Chd4 targets were significantly enriched for both up and downregulated DEG (**Supplementary Figure 8B**, p-value=1.88e-3 and 2.91e-3 for up and downregulated DEG, respectively), supporting dual directionality of modulation by these complexes. However, the presence/absence of ADNP in the POGZ-containing complex showed mirrored effects on gene expression: upregulated DEG were significantly enriched for mouse Pogz/Adnp targets while downregulated DEG were significantly enriched for mouse Pogz targets of complexes without Adnp (**Supplementary Figure 8B**).

Comparison of the complex-informed subsets of POGZ targets with Pogz LoF DEG across different mouse brain regions further emphasized that the direction of modulation of POGZ-regulated genes can vary according to the tissue investigated (**Supplementary Figure 10**). For example, Pogz/Chd4 target genes were significantly enriched among the downregulated mouse cerebellum DEG, yet among the upregulated hippocampus DEG and human compound het iN DEG (**Supplementary Figure 8B, Supplementary Figure 10**). Additional differences in pattern by brain region were evident for other Pogz complex targets indicating that the direction of effect on POGZ direct regulatory targets varies across cell type and possibly species, presenting a complexity that will likely require future single-cell analyses to resolve.

### POGZ haploinsufficiency impacts gene regulation of non-POGZ targets

In some instances, transcription factors (TFs) can produce nucleosome repositioning through the recruitment of chromatin remodelers to promote changes in DNA accessibility. To address a POGZ regulatory impact on chromatin accessibility in the disease-relevant genotypic context, we examined ATAC-seq profiles in the heterozygous cell models.

We first identified differential accessibility regions (DARs) comparing WT and heterozygous deletion lines (**Supplementary Table 8)**, then investigated DARs enrichment among promoters of DEG. We discovered that DEG in both iN and NSC displayed a strong association with DARs in these neuronal models, emphasizing the value of such multiomic profiling (**Supplementary Figure 11**, p-value=9.11e-10 and 7.14e-3 for iN and NSC, respectively). In heterozygous lines, there was no association between POGZ target DEG and chromatin accessibility changes at gene promoters, but we did observe strong association in homozygous models (**Supplemental material**). Additionally, most of the DEG observed in the context of POGZ haploinsufficiency were not genes known to be direct POGZ targets (**Supplementary Figure 6A**). This evidence suggests that DARs and DEG in heterozygous lines might be driven by indirect regulatory mechanisms. Therefore, we defined POGZ’s indirect effects as the impact of other TFs over DEG in heterozygous deletion lines. We hypothesize that these indirect effects might occur due to differential expression of other TFs themselves or due to differential accessibility where other TFs bind (see definitions of indirect effects on **Supplementary Figure 5B**). We then leveraged another layer of ATAC-seq data to explore this hypothesis.

### Indirect effects are mediated by other transcription factors

To gain molecular insight into POGZ’s indirect effects, we profiled accessible binding sites for a broad range of TFs in the haploinsufficiency models. From deep coverage ATAC-seq analysis, we called the “transcription factor footprint”, considered as a proxy for TF binding probability, based on the depletion of ATAC-seq local signal over a TF motif within an open chromatin region^30^ (see Methods for details). The aim of this approach was to reveal those TF whose binding was most susceptible to *POGZ* haploinsufficiency, potentially reflecting indirect POGZ regulation. We dissected the footprints for 429 and 393 motifs, whose corresponding TFs were expressed in iN and NSC models (based on our RNA-seq datasets), respectively (**Supplementary Table 9**). Briefly, for each TF motif we first scored the genome-wide footprint differences between genotypes by summarizing the changes at all of its matching loci. When any motif of a single TF genome-wide change was significant (See Methods), we considered it a differentially footprinted TF (DTF) and further screened its footprint at each motif-containing locus. In iN, the 16 DTFs identified from genome-wide scoring all indicated reduced footprinting in *POGZ* heterozygous LoF cells. Of 39 DTFs in heterozygous NSC, 20 and 19 indicated increased and reduced footprinting, respectively (**Supplementary Table 9**). In iN, 31 (∼2.5%) of 1,245 expressed TFs are DEG and none is a DTF, while in NSC, 89 (∼7.6%) out of 1,168 expressed TFs are DEG and only one, *SNAI2*, is also a DTF. We further confirmed that the potential binding of SNAI2 was not enriched in the promoter of NSC DEG. There were 14 DTFs shared by both cell types and many of these involve related protein complexes. Together they comprise proteins encoded by 11 genes, none of which was a DEG in our heterozygous cellular models (**Supplementary Figure 12A**).

The subset of genes whose promoters contain the DTF binding sites (i.e., DTF targets) overlapped significantly with DEG (**Figure 4A** and **B**). The pathways strongly enriched among differentially expressed POGZ targets (**Supplementary Figure 6C**) were not significantly enriched among differentially expressed DTF targets (**Figure 4C**). Thus, in haploinsufficient *POGZ* models the downregulation of genes related to DNA binding, transcription and RNA processing (**Supplementary Figure 6C**) is specifically caused by POGZ direct regulation not indirect regulatory effects (**Figure 4C**).

**Figure 4.**
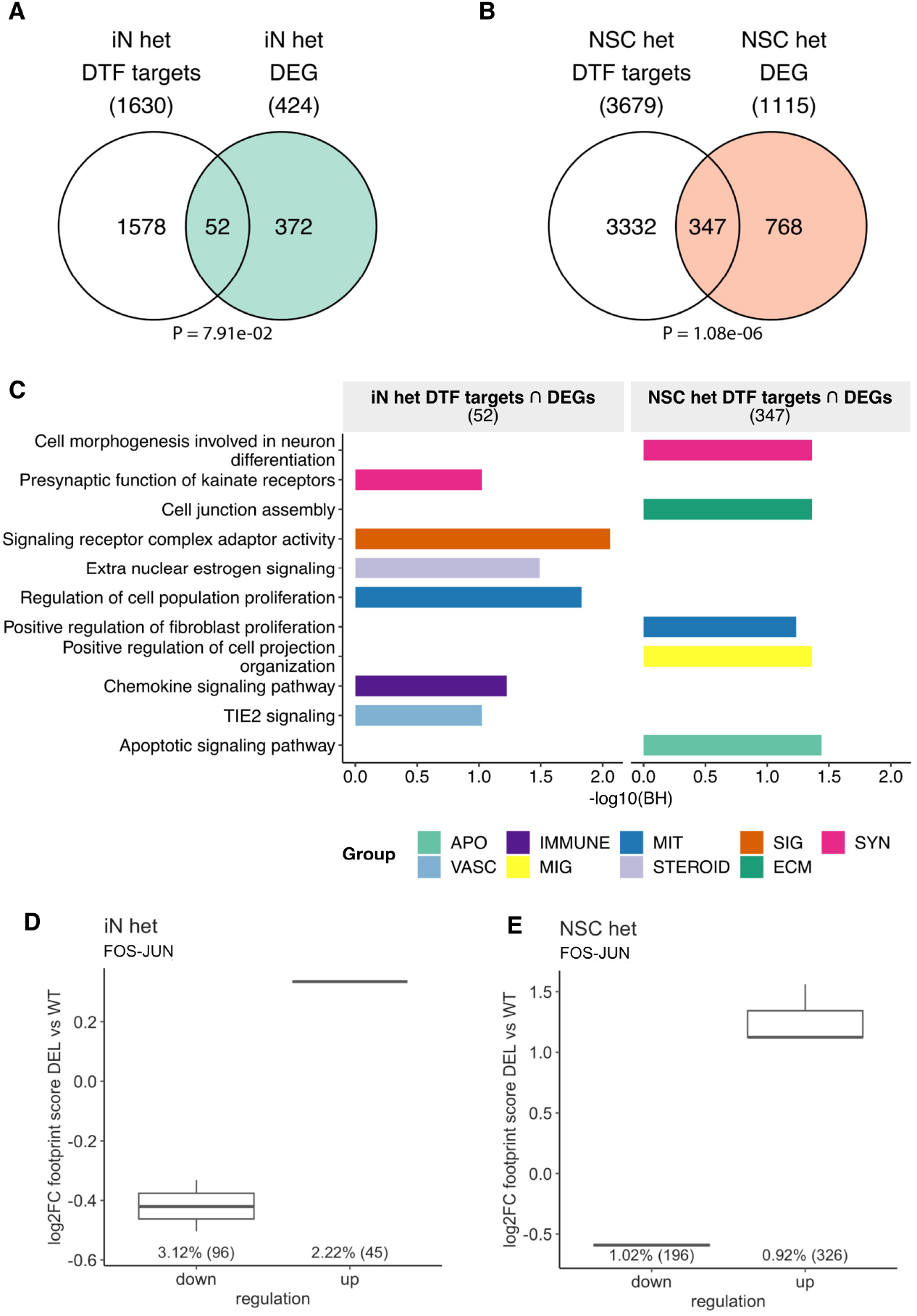
Integration between differential expression and footprinting analysis to identify indirect regulatory mechanisms. **(A)** and **(B)** Overlap between DEG and DTF targets from human heterozygous (het) iN or NSC. **(C)** Statistical significance for enrichment of pathways among heterozygous iN (left panel) and NSC (right panel) DTF targets which are DEG. Intersection of two gene lists is represented by “⋂”. Numbers in parentheses represent total genes in a given list. Colors represent pathway categorization according to the nature of the functional categories related to the enriched terms. Groups of functional categories: APO: apoptosis, IMMUNE: immune system and inflammation, MIG: cell migration, MIT: mitosis and cell proliferation, STEROID: steroid metabolism, SIG: intracellular signaling pathways, ECM: extracellular matrix, SYN: synapse, VASC: vascular. **(D)** and **(E)** FOS-JUN footprinting score over gene promoters of down/upregulated FOS-JUN targets in heterozygous iN (D) or NSC (E) lines. Numbers in parentheses indicate the total number of genes in a given gene list. BH: Benjamini-Hochberg adjusted p-values.

We then sought to ascertain whether any DTF shared between cell types might represent a common element of POGZ regulation. Of 14 DTFs found in both cell types (**Supplementary Table 9**), 13 belong to a well-known immediate early gene transcriptional regulator^31,32^, the AP1 (activator protein-1) complex family, with JUN and/or FOS components (**Supplementary Table 9**). In agreement with the globally increased footprinting of AP1 complexes in NSC and decreased footprinting in iN, AP1 regulatory targets among DEG were mainly upregulated by heterozygous POGZ LoF in NSC and downregulated in iN (**Figure 4D** and **E**). In both cell types, the fold changes of footprinting at the AP1 complex binding sites were in the same direction as the expression of their target genes (**Figure 4D** and **E**), in accordance with the transcriptional activator activity previously described for AP1 complexes^31,32^. Because neither this nor any other class of TFs was enriched among overall DEG, it is the activity of this class of TFs that is modulated by POGZ haploinsufficiency. Their impact largely differs in direction between NSC and iN models, but accounts for a substantial portion of the altered gene expression in NSC and iN.

### Neuronal haploinsufficiency of two NDD-associated chromatin genes shows molecular pathway convergence

Chromatin remodelers and transcriptional regulators constitute functional classes consistently found to be disrupted in large-scale exome sequencing studies of ASD/NDD^1,7,13^. To identify shared molecular signatures by which chromatin-related NDD-risk genes might contribute to pathology, we compared the transcriptional profiles from independent neuronal haploinsufficiency models for *POGZ* and *MEF2C*, another well-established NDD-associated gene, encoding Myocyte Enhancer Factor 2C^33^. *MEF2C* was also found to be a DTF target in the *POGZ* heterozygous deletion NSC.

The hiPSC-derived neural models (iN and NSC) for both gene perturbations were generated strictly using the same CRISPR editing and differentiation protocols^33^. Since the *MEF2C* models were created on the GM08830 hiPSC background, we focused on the *POGZ* GM08830 iN and NSC cells, assuring a direct isogenic comparison of the RNA-seq datasets not only between conditions (i.e., edited vs. unedited clones) but also between genes (i.e., *POGZ* vs. *MEF2C*). There was a significant overlap of DEG at FDR<0.1 between these haploinsufficiency models (**Figure 5A**). Shared DEG in iN were significantly enriched for processes related to intracellular signaling and extracellular matrix organization (**Figure 5C**), while in NSC, they were related to chromatin and transcription regulation, including gene targets of AP-1 complexes (**Supplementary Figure 13**).

**Figure 5.**
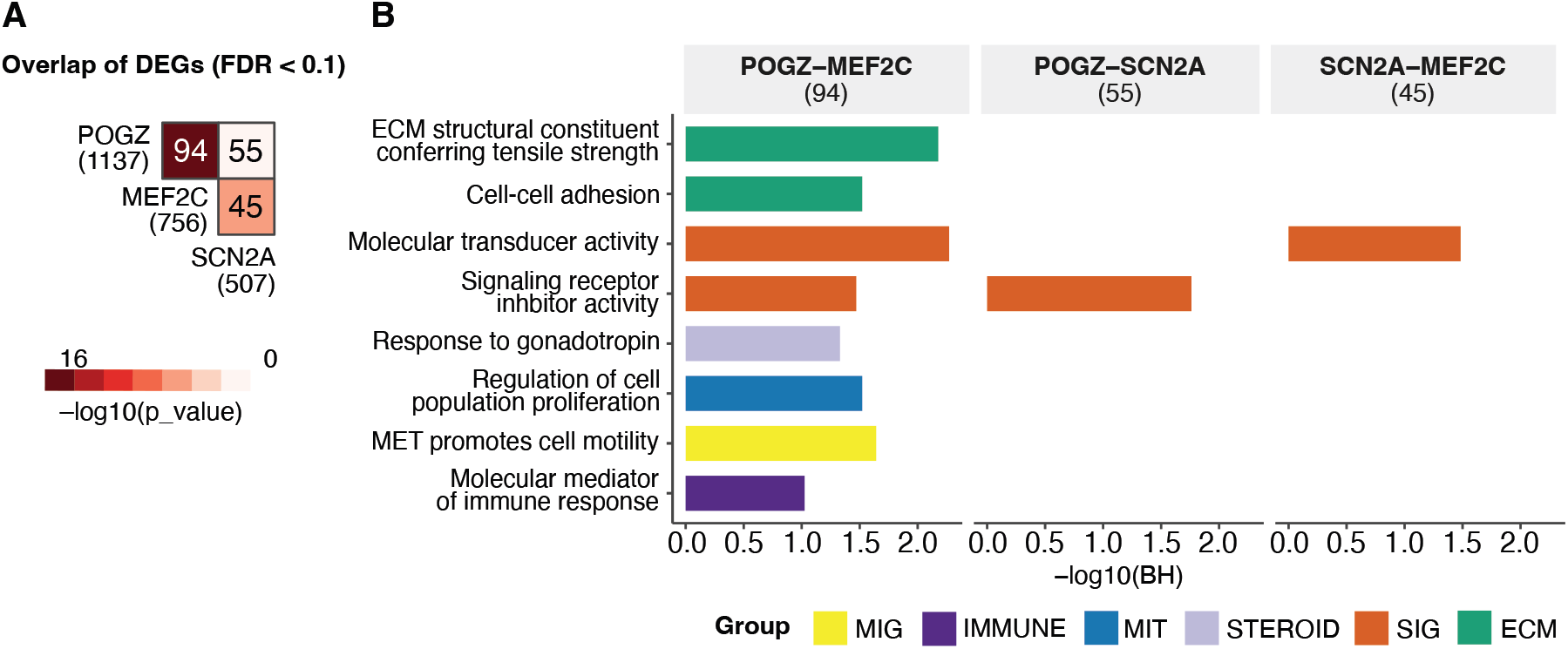
Overlap of transcriptional profiles derived from isogenic heterozygous iN with POGZ, MEF2C or SCN2A gene deletions. **(A)** Heatmap indicating statistical significance for overlap between DEG (FDR<0.1) from heterozygous iN with POGZ, MEF2C, or SCN2A gene deletions. **(B)** Statistical significance for enrichment of pathways among DEG overlaps from two-way comparisons of heterozygous iN with POGZ, MEF2C or SCN2A gene deletions. Numbers in parentheses indicate the total number of genes in a given gene list. Colors represent pathway categorization according to the nature of the functional categories related to the enriched term. Groups of functional categories: IMMUNE: immune system and inflammation, MIG: cell migration, MIT: mitosis and cell proliferation, STEROID: steroid metabolism, SIG: intracellular signaling pathways, ECM: extracellular matrix. BH: Benjamini-Hochberg adjusted p-values. Numbers in parentheses indicate the total number of genes in a given gene list.

### Similarities and differences due to haploinsufficiency of NDD-associated chromatin and synapse genes in human neurons

We next expanded our functional analyses to compare our *POGZ* LoF models with LoF of another NDD-associated gene with a different biological role: *SCN2A* encoding the alpha 2 subunit of the voltage-gated sodium channel, thought to cause NDD through its effect on synaptic activity, is among the most robustly associated ASD genes^1^. Since *SCN2A* is expressed at very low levels in NSC, we only focused this comparison on iN edited lines generated on the same isogenic background as those for *POGZ* using similar procedures.

We performed WGCNA co-expression analysis considering heterozygous mutant iN models. Both heterozygous deletion and control iN samples for *POGZ* (23 samples) and *SCN2A* (35 samples) from GM80830 background were considered, whereas *MEF2C* iN samples (11 samples) were excluded as they did not meet the criteria for the minimum number of clones (see Supplemental Information for details).

The “green” module (N=647) presented discordant regulation between *POGZ* and *SCN2A* edited lines (upregulated and downregulated, respectively **Supplementary Table 10**). It was strongly enriched for ASD/NDD associated genes and LoF constrained genes and mildly enriched for chromatin modifiers, but not synaptic genes (**Supplementary Figure 14, Supplementary Figure 15**). This suggested that *POGZ* and *SCN2A* haploinsufficiency can impact some of the same neurodevelopmental processes, but in opposite directions. The “tan” (N=395) and “purple” (N=603) modules also reflected opposing transcriptional effects but without disease-associated enrichments. Only in the “midnightblue” module (N=341) was the direction of effect the same (downregulation) for both *POGZ* and *SCN2A* haploinsufficiency. Together, these analyses suggested a mild convergent signal between *POGZ* and *SCN2A* haploinsufficiency involving co-modulation of NDD-associated and constraint genes, but with transcriptional changes in opposite directions.

We next employed microelectrode array (MEA) for electrophysiological comparison of the isogenic iN with heterozygous *POGZ* or *SCN2A* gene deletions (**Figure 1D**). The mild WGCNA-defined differences did not translate into alterations in firing rate, where subtle effects did not achieve statistical significance. In contrast, when both alleles were edited to achieve complete LoF, both *POGZ* and *SCN2A* models displayed dramatic effects on neuronal firing rate, but in opposite directions, with increased firing in *POGZ* compound heterozygous iN and decreased firing in homozygous *SCN2A* iN (**Figure 6A** and **B**). Changes in neurite length, measured using label-free Incucyte, confirmed the strong impact of the total functional inactivation of these genes in iN (**Figure 6C** and **D**), with *POGZ* compound heterozygous iN exhibiting a more intricate neuronal network and homozygous *SCN2A* lines showing reduced neurite length and branching (**Figure 6**).

**Figure 6.**
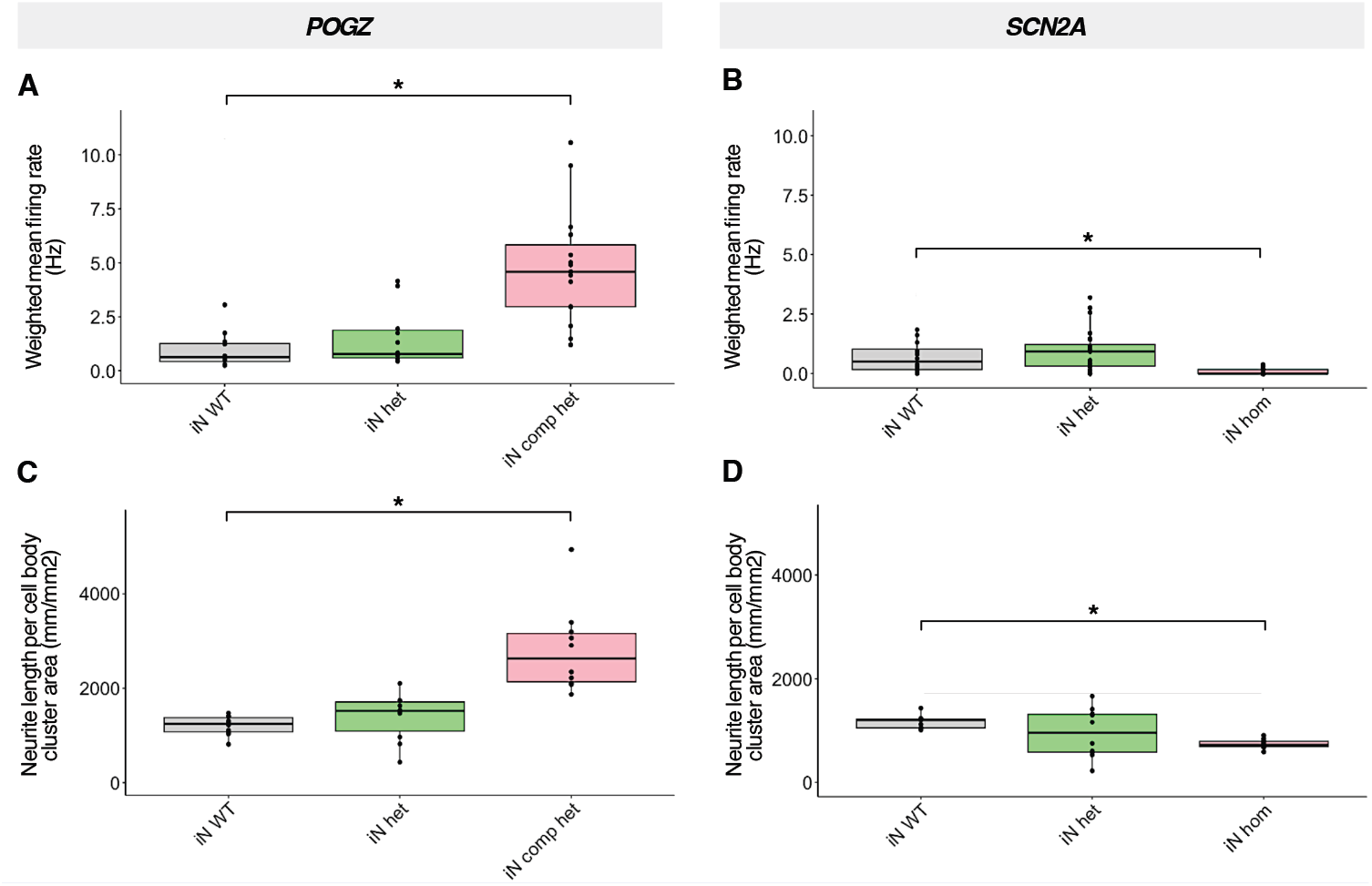
Cellular phenotyping on isogenic heterozygous iN with POGZ or SCN2A gene deletions. **(A)** and **(B)**. Weighted mean firing rate at the time point with highest activity measured by multi-electrode array in iN with POGZ heterozygous (het) and compound heterozygous (comp het) deletions (A) and with SCN2A het and homozygous (hom) deletions (B). **(C)** and **(D)**. Neurite length normalized by cell body cluster area at the time point with highest extension density measured by Incucyte in iN with POGZ heterozygous (het) and compound heterozygous (comp het) deletions (C) and with SCN2A het and homozygous (hom) deletions (D). Asterisk indicates p-value<0.01.

In conclusion, while heterozygous LoF of *POGZ* and *SCN2A* exhibited subtle and often opposing transcriptional effects, particularly within modules that were significantly enriched for NDD-associated genes, these changes did not translate to significant alterations in neuronal firing rate or neurite extension in iN. However, the complete, biallelic LoF of these genes revealed robust yet contrasting impacts on neuronal function, arguing that while both genes contribute to NDD their specific mechanisms and the direction of their impact might differ.

## DISCUSSION

Unlike functional studies using cell lines derived from patients, introducing genomic variants of interest into an isogenic background permits direct comparisons across genes and mutations for a broader scope of evaluation of regulatory mechanisms, including efforts to infer causal relationships between variant, expression profiles, chromatin accessibility regulation, and neuronal phenotypes^34–36^. Our CRISPR engineering experiments yielded a catalog of 66 hiPSC replicates, with different zygosities across two different background lines. The integration of sequencing datasets from this compendium of models allowed us to statistically dissect the molecular signals which are consistent between multiple experiments. The significant overlap between the biological signatures observed in models derived from different hiPSC backgrounds indicated that the impacts due to *POGZ* LoF described here are reproducible.

The comparison across different cell types allowed a broader scope of evaluation of regulatory mechanisms. It has been demonstrated that the expression of chromatin-related NDD genes peaks in early neurodevelopment (8-9 human gestational weeks)^37,38^ and, when detected in later stages, is enriched in cortical layers, including excitatory neuronal lineages^39^. Additionally, broader genetic studies in neuropsychiatric diseases^40^ and single-cell transcriptomics in ASD^41^ have implicated neuronal genes that converge on cellular programs associated with excitatory neurons. Thus, the targeted cell types investigated here – NSC and iN – correspond to disease-relevant contexts at early developmental stages and mature synaptic functioning, respectively. Functional impacts due to *POGZ* LoF frequently affected similar gene sets in NSC and iN; however, in many instances, the directions of effect were opposite in the two cell types. These differences were partially related to the specific composition of POGZ protein complexes, and the gene targets directly regulated by those different versions.

In agreement with previous *POGZ* functional investigations^22^, *POGZ* heterozygous models presented with subtle molecular impacts, especially regarding transcriptional disturbances of POGZ target genes. In heterozygous models, these transcriptional disturbances are partially derived from indirect regulatory effects (i.e, transcriptional disturbance of DTF target genes), and AP1 complexes are the main class of TFs with regulatory function affected in both cell types. AP1 complexes are long-standing immediate early gene regulators, with drastic implications for synaptic activity^42^. Mouse knockouts for *Actl6B*, a BAF complex component, have previously shown impairment in early gene response through disruption of AP1 complex regulatory activity^43^. Additionally, increased c-fos protein expression associated with higher excitatory synaptic inputs have been observed in the mouse brain with the LoF of *Auts2*, a well-known ASD-associated gene^44^.

Beyond *POGZ*-specific functional impacts, this study took advantage of CRISPR-engineered isogenic neuronal heterozygous models to explore the molecular convergence between *POGZ* and other ASD/ NDD benchmark genes from different functional classes: *SCN2A*, the top ASD-associated synaptic gene, and *MEF2C*, which is one of the main NDD-associated TFs and was also found to be a DTF target in lines with *POGZ* heterozygous deletion. *POGZ* and *MEF2C* showed strong convergence of their molecular impacts in NSC and iN, with substantial overlaps on overall transcriptional disturbance and affected biological pathways, including disruption of AP1 complexes regulation and extracellular matrix. Previous MEA studies performed by our research group applied analogous functional protocols on isogenic neuronal lines with *MEF2C* LoF^33^. When the results from this study are compared to the MEA data presented here, we can conclude that LoF of POGZ and MEF2C show opposite effects on firing activity, with *POGZ* LoF models showing increased firing rate, and the isogenic models of MEF2C LoF showing decreased firing rate. These results suggest that despite the striking transcriptional convergence between *MEF2C* and *POGZ* iN lines, shared transcriptional signals between different NDD/ASD genes might be translated into effects on the same cellular processes, but in different directions.

Co-expression networks in heterozygous iN showed that constrained NDD-associated genes were modulated in opposite direction in *POGZ* and *SCN2A* models, with inverse functional consequences in the cellular assessments between these gene perturbations. The decrease in neuronal excitability in human iN with biallelic mutations in *SCN2A* is concordant with previous studies ^34^, while the increased excitability in POGZ models for POGZ biallelic mutations is unprecedented in the literature and might be specific for the cell type here evaluated. Although the complete removal of *POGZ* or *SCN2A* do not reflect the condition in human disease, the biallelic models generated in this study provided mechanistic insights associated with neuronal abnormalities. Alterations in the excitation-inhibition balance ratio in cerebral cortex is a common feature in animal models of rare genetic variants associated with ASD and have been proposed as a relevant pathophysiological mechanism in ASD^45–47^. The altered excitability observed in our knockout *POGZ* and *SCN2A* models might be relevant to understanding the cellular consequences of these genomic perturbations in the in vitro neuronal used in this study.

Our study design presents some limitations. Monolayer cellular cultures fail to capture the cellular complexity of the human brain. Although NSC and iN can be considered as representative of different developmental stages, they are not appropriate models for trajectories of cellular differentiation. This study considered POGZ direct regulatory targets defined in previously published functional experiments which were based on human fetal cortex and mESC^22,23^. POGZ binding sites in iN and NSC might differ; however, the overall profile is expected to be conserved across cell types, given the strong overlap observed between human fetal cortex and mESC POGZ targets. These limitations indicate that our results should be interpreted cautiously with respect to clinical implications. It is important to note that more than half of DEG were neither POGZ known targets nor DTF targets, and other regulatory mechanisms remain to be identified. Nevertheless, by addressing the effect of the interrogated gene disruptions on specific neuronal cell types with isogenic backgrounds, we were able to dissect the functional assessment of isolated cell populations of great interest in NDD pathogenesis, resulting in unambiguous data interpretation.

We present CRISPR-engineered isogenic allelic series of *POGZ* disruption in human-derived in vitro models and functional characterization of resultant transcriptional and chromatin accessibility effects. These studies uniquely dissect the parallel consequences of *POGZ* direct and indirect regulatory effects on heterozygous cell lines, which carry the disease-relevant genotypic context. Our study clearly demonstrates convergence between ASD/ NDD genes and highlights commonly affected pathways, which might be pharmacologically targeted in future investigations.

## METHODS

### Engineering of CRISPR-mediated gene deletions

Human induced pluripotent stem cell (hiPSC) lines were maintained on Matrigel-coated dishes (Corning) with Essential 8 culture medium (Thermo Fisher Scientific) and incubated at 37°C in a humidified atmosphere with 5% CO2. CRISPR manipulations on *POGZ* locus were performed in two hiPSC lines with independent genetic backgrounds, using one male (GM08830^48^) and one female line (MGH2069^49^). Those two hiPSC lines were previously deep whole genome sequenced to confirm the absence of NDD-associated variants. Characterization of pluripotency has been previously described^50^. Two *POGZ* deletion types (one full and one partial gene deletion) were attempted to increase the probability of editing efficiency. Both deletion types are intended to cause POGZ LoF. Cas9 guide RNA (gRNA) sequences were designed immediately 5′ and 3′ to POGZ gene and at POGZ intron 12 (NM_015100, gRNA sequences are available in **Supplementary Table 11**) and obtained as modified synthetic sgRNA (Synthego). hiPSCs were co-transfected with the ribonucleoprotein (RNP) complex [Cas9 (Alt-R® S.p. HiFi Cas9 Nuclease V3, Integrated DNA Technologies)+gRNA] and CleanCap EGFP mRNA (TriLink Biotechnologies) using Lipofectamine Stem Transfection Reagent (Invitrogen) according to manufacturer’s instructions. To generate the whole gene deletion (del1, 56kb) and partial gene deletion (del2, 7kb) in heterozygous state, wild type hiPSCs were transfected with two different combinations of gRNAs: (1) 5’ plus 3’ gRNAs and (2) intron 12 plus 3’ gRNAs (**Figure S1A** and **S1B**). To generate the compound heterozygous lines for *POGZ* locus, hiPSCs with POGZ whole gene deletion (del1) were transfected with intron 12 plus 3’ gRNAs (**Figure S1A** and **S1B**). gRNAs and target editions for CRISPR engineering of *SNC2A* and *MEF2C* gene deletions were previously described^33,51^. 48 h after transfection, cells were separated by fluorescence-activated cell sorting (FACS) and during the first four days after FACS, hiPSCs are cultured in Essential 8 medium supplemented with CloneR (StemCell Technologies), as previously described ^52^. In the CRISPR edition experiment to generate POGZ compound heterozygote lines, a wild type hiPSC line was transfected with Cas9 and GFP, subjected to FACS, and the resultant clones were used as wild type controls in the subsequent experiments. Once individual colonies were available, they were split into two populations, one for DNA extraction and genotyping (see section below) and one for future propagation. Successfully edited, clonal colonies were expanded as well as CRISPR-exposed unedited clones from the same transfection and sorting experiment, which were used as controls in downstream analysis. All replicates of edited and control clones were independently submitted to TRA-1-60 sorting, using anti-TRA-1-60 microbeads (Miltenyi Biotec) to select pluripotent cells and only selected cells were used in subsequent differentiation procedures.

### Characterization of CRISPR-mediated edits

Prior to clone expansion, genomic DNA from individual hiPSC clones was isolated using Quick-DNA 96 kit (Zymo Research). Each clone was genotyped using PCR followed by Sanger sequencing to identify colonies that acquired desired deletions (**Supplementary Figure 1A** and **1B**, primer sequences are described in **Supplementary Table 11**). 16 out of 22 deletions’ junction points presented with insertions and indels involving few base pairs, ranging from 1 to 26Bp, none of them affecting coding regions or splice sites. To ensure clonality and determine gene dosage, each clone was genotyped by digital droplet PCR (Bio-Rad) using *POGZ* TaqMan copy number probe (Thermo Fisher Scientific, assay Hs00213580_cn), and Rnase P TaqMan Copy Number Reference Assay (Thermo Fisher Scientific) for normalization. This clone genotyping information and total clone survival rate after FACS was used to calculate the editing efficiency for each condition (**Supplementary Table 12**). All successfully edited clones, and an equal number of control clones (i.e. unedited CRISPR-exposed clones from the same FACS batch) were expanded for future functional analysis. During clone expansion, four passages after FACS, genomic DNA from each edited and control clone was re-extracted using DNeasy Blood & Tissue Kit (Qiagen) and copy number state was re-evaluated using the same ddPCR assays (**Supplementary Figure 1C**). At this passage, the CRISPR cut sites in wild type alleles from heterozygous edited clones and control clones were assessed by PCR and sanger sequencing (primer sequences are described in **Supplementary Table 11**). Two clones with heterozygous deletions presented with a 6Bp indel and a 1Bp insertion at intron 12 and 3’ to POGZ cut sites, respectively, and none of these events affected coding regions or splice sites. All control clones used in subsequent experiments did not show insertions or indels at Cas9 cut sites. Genotyping of *SNC2A* and *MEF2C* gene deletions were previously described^33,51^.

### Replicates and batches design for neuronal differentiation experiments with POGZ models

Differentiation of *POGZ* heterozygous hiPSC lines into neuronal lines was conducted on batches of six CRISPR/Cas9 edited replicates and six control replicates per *POGZ* deletion per background line, resulting in four batches of 12 clones in total (**Supplementary Table 13**). Differentiation of *POGZ* compound heterozygous hiPSC lines into neuronal lines was conducted on one batch of six compound heterozygous replicates, six heterozygous whole gene deletion replicates and six wild type control replicates (**Supplementary Table 13**). Independently edited clones were used as replicates in the following experiments when possible. For wild type hiPSCs CRISPR transfection/FACS experiments that generated less than six clones with heterozygous deletions, one or two edited clones were split during expansion to reach the total of six replicates (**Supplementary Table 13**). Since CRISPR transfection/ FACS experiments to generate compound heterozygote lines generated a single edited clone, this one clone was split into six replicates for future experiments (**Supplementary Table 13**). After neuronal differentiation, given a certain cell type, the replicates for RNA-seq and ATAC-seq experiments were harvested in parallel.

### Neural induction of hiPSC lines and expansion of NSC

Neural induction of hiPSC lines was performed with PSC Neural Induction Medium (Thermo Fisher Scientific) according to the manufacturer’s instructions. hiPSC clones were at 5-8 passages after CRISPR transfection when neural induction was initiated. Briefly, on day 0 of neural induction, ∼24 h after the cells were split, the Essential 8 medium was replaced with PSC neural induction medium, which contains neurobasal medium and PSC neural induction supplement. The neural induction medium was exchanged every other day from day 0 to day 5 of neural induction. On day 7, NSC (P0) were harvested and expanded in neural expansion medium containing 50% neurobasal medium (Thermo Fisher Scientific), 50% advanced DMEM/F12 (Thermo Fisher Scientific), and neural induction supplement (Thermo Fisher Scientific) on Matrigel. At passage 4, all clones were submitted to NCAM sorting using anti-PSA-NCAM microbeads (Miltenyi Biotec) to select neural cells and only the selected cells were plated at passage 5. At passage 7, NSC were plated on coverslips and analyzed for Nestin, PAX6, SOX1 and SOX2 expression by immunostaining using Human Neural Stem Cell Immunocytochemistry Kit (Thermo Fisher Scientific) according to manufacturer’s instructions (**Supplementary Figure 16**). Expanded NSC at passage 7 were harvested and used for subsequent assays.

### Validation of POGZ expression reduction

After neural differentiation, the impact of the edits on transcript and protein abundance was measured by digital droplet PCR (Bio-Rad) and western blot, respectively. For digital droplet PCR gene expression assays, RNA was isolated with Trizol reagent following manufacturer’s instructions, and cDNA was synthesized using Maxima H Minus cDNA Synthesis kit (Thermo Fisher Scientific). POGZ transcript abundance was accessed with TaqMan gene expression probes (Thermo Fisher Scientific, assays Hs00374850_ m1 and Hs00418559_m1) and normalized by GUSB gene expression (Hs00939627_m1). For western blot assays, fractionated protein extraction was performed using NE PER Nuclear and Cytoplasmic Extraction Kit (Thermo Fisher Scientific). Nuclear extracts were resolved on NuPAGE™ 3-8% Tris-Acetate Gels (Thermo Fisher Scientific) and wet-transferred to PVDF membrane (Millipore Sigma). Membranes were blocked with Odyssey Blocking Buffer (LI-COR Bioscience) for 60 minutes at room temperature and then probed overnight at 4°C with anti-POGZ (Abcam ab171934, 1:300) and anti-PCNA (Abcam ab29, 1:1000), which was used for normalization. After three washes, membranes were incubated with IRDye secondary antibodies (LI-COR Bioscience) (**Supplementary Figure 1D**).

### Induction of Ngn2-neurons from hiPSC lines

Ngn2-neuronal induction was performed as previously described^53^ with modifications. Briefly, hiPSCs were seeded at low density and transduced with a lentivirus expressing TetO-Ngn2-GFP-Puro or TetO-Ngn2-Puro along with rtTA. Twenty-four hours after transduction, doxycycline was added to initiate Ngn2 expression, and then 24 h later the cells were selected with puromycin. hiPSC-derived *Ngn2*-induced neurons were cultured in neuronal maintenance medium supplemented with BDNF and NT-3 growth factors. Subsequent experiments were performed with 3-week-old *Ngn2*-induced neurons.

### RNA-seq on neuronal models

RNA was isolated with Trizol reagent (Thermo Fisher Scientific) and Phasemaker Tubes (Invitrogen) following manufacturer’s instructions. Both *POGZ* and *SCN2A* RNA-seq libraries were prepared using 200 ng of total RNA using a TruSeq Stranded mRNA Sample Prep Kit (Illumina cat # RS-122-2102). Libraries were multiplexed, pooled and sequenced on an Illumina NovaSeq, generating an average of 40.7 million paired-end 150-cycle reads for 133 *POGZ* samples and an average of 33.5 million paired-end 150-cycle reads for 38 *SCN2A* samples.

### RNA-seq data analysis

Quality of sequence reads was assessed by fastQC (version 0.10.1) (https://www.bioinformatics.babraham.ac.uk/projects/fastqc/). Raw sequence reads were trimmed against Illumina adapters using Trimmomatic (v. 0.36) with parameters ILLUMINACLIP:adapter. fa:2:30:10 LEADING:3 TRAILING:3 SLIDINGWINDOW:4:15 MINLEN:75 (https://www.ncbi.nlm.nih.gov/pmc/articles/PMC4103590/). We generated gene-based counts for RNA-seq libraries by aligning sequence reads to the human reference genome, GRCh38 (v92), and relying on Ensembl gene annotations of this reference genome by using STAR (version 2.5.2A)^54^ with parameters ‘‘–outSAMunmapped Within –outFilterMultimapNmax 1 –outFilterMismatchNoverLmax 0.1 –alignIntronMin 21 – alignIntronMax 0 –alignEndsType Local –quantMode GeneCounts –twopassMode Basic’’. We assessed the quality of alignments by using custom scripts in Picard Tools, RNA-seqC^55^, and SamTools^56^. These quality-checking assessments and exploratory analyses identified two outlier samples from POGZ samples: one NSC del2 sample from MGH2069 background (NSC_03_2020_KOK25_14) and one iN WT sample from GM08830 background (iN_05_2021_ 8330_WT_C6), and three outlier samples from SCN2A samples: two edited samples with SCN2A heterozygous deletion (CD1-55, CD3-50) and one WT sample (CD3-43).

To validate the deletion introduced by CRISPR on *POGZ* and *SCN2A* loci, raw exon-level coverage counts was generated using DEXseq^57^ on all annotated transcript isoforms (**Supplementary Tables 14 and 15**). Raw exon-coverage counts were further normalized by exon length and number of uniquely mapped reads which were retrieved from STAR aligner’s log file for each library, following the formula 10^9^ x raw_count_exon_ij_ / (Uniquely Mapped Reads_j_ x Exon Length_i_) where index i denotes the exon number in the j^th^ library. For each library, normalized coverage of deletion region was statistically compared to normalized coverage of exons outside the deletion region applying one-tailed t-test.

In each comparison of target gene deletion vs WT samples, genes with more than 0.5 counts per million (CPM) in at least 50% of samples in either condition were analyzed. In gene-level CPM calculations, the number of uniquely mapped reads output from STAR aligner was used as library size.

### Differential expression analysis

Differential expression analysis for *POGZ* iN (heterozygous deletion vs WT) and *SCN2A* iN (heterozygous deletion vs WT) RNA-seq libraries were performed using DESeq2^58^ + SVA^59^ pipeline in which surrogate variables were estimated using R/Biocondutor package, SVA with ∼genotype as the full model and ∼1 as reduced model. In differential expression analysis of *POGZ* iN samples, del1 and del2 genotypes were combined under del for each background, and DESeq2 + SVA pipeline was applied to each background. Shared genes that were significant at FDR<0.1 and showed the same direction of effect in both backgrounds comprised iN DEG. NSC DEG were identified similarly as shared genes that were significant at FDR<0.1 and showed the same direction of effect in both backgrounds. However, differential expression of genes in each background was assessed by integrating del1 and del2 through meta-analysis. In doing so, in each background, differential expression of each gene due to del1 and del2 was assessed by z scores. These z scores were combined under Stouffer’s weighted z score method and converted to p-value following normal distribution. Adjusted p-value (FDR) were calculated separately for each background, following Benjamini-Hochberg procedure. In each background, DEG were defined among genes that were significant at FDR<0.1 and showed the same direction of effect due to del1 and del2. Only DESeq2 was used for differential expression analyses of POGZ (compound heterozygous vs WT) in NSC and iN as principal component analysis revealed that compound heterozygous samples and control samples were distinctly clustered.

### Co-expression analysis

Co-expression network analysis was performed using R package WGCNA ^28^ for each cell type separately using the signed network type for which we used log transformed CPM filtered and SVA corrected counts. Soft power was selected such that the scale-free topology fit (R2)>0.85. and the smallest module size was set to 50. Merge of the modules with similar eigengene profiles was performed (Similarity>75%). Module membership for each gene was re-evaluated based on the module membership p-value; genes with p-value>0.01 were marked as unassigned (gray module 0). Modules of interest in NSC and iN were identified based on statistical significance of module eigengene correlation with *POGZ* genotype. In co-expression analysis of chromatin and synaptic ASD/NDD genes, both heterozygous deletion and control iN samples for *POGZ* (23 samples) and *SCN2A* (35 samples) from GM8083 background were considered, whereas *MEF2C* iN samples (11 samples) were excluded as they were fewer compared to others. SVA corrected counts were generated by applying SVA package to counts that were normalized by DESeq2 under the full model ∼ gene_genotype, where gene_genotype consists of three groups of genotype; *POGZ* heterozygous deletion combining both del1 and del2, *SCN2A* heterozygous deletion and control samples combining control samples from *POGZ* del1 and del2 batches, and *SCN2A* batch and ∼1 as reduced model. One *SCN2A* WT sample (CD3-38) was identified as an outlier according to previously described procedures^60,61^

### Enrichment analysis

We tested gene enrichments (e.g. DEG and co-expression modules) of DEG and co-expression module genes using one-tailed Fisher’s exact test for the curated lists of Gene Ontology (GO) Terms, Canonical Pathways including REACTOME and KEGG, Human Phenotype Ontology terms, and TF targets from mSigDB (version 2022.1.Hs)^62,63^, and synaptic-function associated GO terms from SynGO (v1.1)^64^. Additionally, we used phenotype-informed literature data including gene sets previously published with functional associations, mutational constraints or with neurological phenotype, and *POGZ* function. Gene sets with functional associations and mutational constraints are chromatin modifiers and synaptic genes from SynGO (v1.1)^64^, chromatin modifiers^65^, mRNA targets of fragile X messenger ribonucleoprotein (FMRP targets)^66^, and loss-of-function (LoF)-constrained genes^8^. Gene sets associated with neurological phenotype are ASD-risk genes ^1^ and NDD-risk genes^3^. Gene sets related to *POGZ* function are DEG from multiple brain tissues of *Pogz*-deficient mouse models including DEG from mouse hippocampus and cerebellum^21^, and DEG from mouse basal ganglia and cortex^22^, as well as mouse Pogz targets^23^ and human *POGZ* targets^22^. The resulting p-value were corrected for multiple tests using Benjamini-Hochberg correction^67^.

### ATAC-seq on neuronal cell models

ATAC-seq was applied to NSC and iN with the following genotypes: six compound heterozygous clones, six heterozygous whole gene deletion clones and six wild type control clones. ATAC-seq was performed as previously described^68^ with minor modifications. Briefly, 50,000 cells were washed with ice-cold PBS, then re-suspended in 50ul cold Lysis Buffer [10 mM Tris-HCl (pH 7.5), 10 mM NaCl, 3 mM MgCl2, 0.1% (vol/vol) IGEPAL, 0.1% (vol/vol) Tween-20, 0.01% (vol/vol) Digitonin] and kept on ice for 3 minutes. 1 ml Wash Buffer [10 mM Tris-HCl(pH 7.5), 10 mM NaCl, 3 mM MgCl2, 0.1% (vol/vol) Tween-20] was added and samples were centrifuged for 10 minutes at 4°C and 500g. After lysis, the nuclei were re-suspended in 50 ul transposition reaction mix using reagents from Tagment DNA TDE1 Enzyme and Buffer Kits (Illumina) [2X Tagment DNA Buffer, 0.1% (vol/vol) Tween-20, 0.01% (vol/vol) Digitonin, 2.5 μl Tn5 Transposase] and incubated for 30 min at 37°C with constant agitation at 1000rpm. The transposed DNA was purified with MinElute Reaction Cleanup kit (Qiagen) and eluted in 20ul Elution Buffer. DNA fragments were PCR pre-amplified using NEBNext High-Fidelity 2X PCR Master Mix (New England Biolabs) for 5 cycles initially, and one tenth of the volume was removed for qPCR amplification for 20 cycles. A plot of R vs cycle number was generated, and the number of cycles required to reach one third of the maximum R was determined for each sample. The pre-amplified ATAC-seq libraries were then amplified for the calculated additional cycles. Agencourt AMPure XP beads (Beckman Coulter) were used for size selection according to manufacturer’s instructions with 1.8:1 beads to DNA ratio. The size-selected libraries were run on the TapeStation D1000 tape and reagents (Agilent) in order to determine fragment size profile. Libraries were multiplexed, pooled and sequenced on an Illumina NovaSeq 6000, generating an average of 50 million paired-end 50-cycle reads for each sample.

### ATAC-seq analysis

#### Data processing

ATAC-seq data were first processed based on their sequencing batches. For each batch, the processing followed standard ENCODE ATAC-seq workflow (https://docs.google.com/document/d/1f0Cm4vRyDQDu0bMehHD7P7KOMxTOP-HiNoIvL1VcBt8/edit). Briefly, for each library within a batch, the adapters were trimmed, and the reads were aligned to the GRCh38 human genome assembly including mitochondria using Bowtie2 (v2.3.1). Post-alignment quality control approaches were applied, including investigating mitochondria fraction, GC bias, fragment length, phantom peaks and removing PCR duplicates using customized scripts and PicardTools (v1.128). The BAM file was further converted to TagAlign format, where the coordinates adjustment due to Tn5 cleaves was applied by shifting the start positions four bases toward the downstream (i.e. +4 for reads aligned to the positive strand and -5 for reads aligned to the negative strand in a TagAlign file). This TagAlign file was then shuffled and randomly splitted into two files with equal number of lines, which made the two pseudo replicates of that library (i.e. pseudo replicate 1 and 2). For libraries with the same genotype in a batch, they were considered as biological replicates. For each genotype, all the biological replicates, all the pseudo replicate 1’s and all the pseudo replicate 2’s were pooled respectively, generating three extra artificial libraries. Peak calling was then carried out within each genotype on all the libraries, their pseudo replicates and the three pooled artificial libraries, using MACS2^69^ version 2.2.7.1 with “-g hs -p 0.01 --nomodel --shift -75 --extsize 150 --keep-dup all -B –SPMR --call-summits” options. This adds up to 3*(N+1) calls for a genotype with N libraries sequenced. Only the top 300,000 peaks with the smallest p-value from each call were retained for the downstream analyses. To find consensus peaks in each genotype across all the biological replicates, pseudo replicates and artificially pooled libraries, a naive overlap approach was applied, using an overlap cutoff of 50%. This approach searched for overlapped peaks between any two biological replicates that further overlapped with the peaks from the artificially pooled library (i.e. the artificial library where all the biological replicates were pooled); and (b) the overlapped peaks between the two pooled pseudo replicates that further overlapped with the peaks from the artificially pooled library (i.e. the artificial library where all the biological replicates were pooled). This generates 0.5*N*(N-1)+1 overlap sets for a genotype with N libraries sequenced. Each overlap set also excluded the genomic regions recorded in the ENCODE blacklist (hgdownload. cse.ucsc.edu/goldenPath/hg19/encodeDCC/wgEncodeMapability/). For each genotype within a sequencing batch, the overlap set with the greatest number of surviving peaks in (a) was considered as the set of conservative consensus peaks while the overlap set with the greatest number of surviving peaks in both (a) and (b) was considered as the set of optimal consensus peaks. Note, only the conservative consensus peaks were used throughout this study. Finally, the consensus peaks were merged across sequencing batches and genotypes to generate a merged peak set per cell type per iPSC background.

#### Differential Accessibility

The bioconductor package DiffBind (v3.4.11)^70^ was applied to identify DARs between genotypes within each hiPSC background, using the final alignments of ATAC-seq (i.e. BAM files) and the consensus peaks per genotype per sequencing batch as the input. POGZ compound heterozygous peaks were compared to peaks in wild type clones, using dba.contrast function, model design=“∼Treatment” and contract=c(“Treatment”, “DEL”, “WT”). “Treatment” represents sample genotype (del1, del1_del2, or WT). POGZ heterozygous deletion peaks in GM08830 and MGH2069 were compared to their corresponding peaks in wild type clones, respectively, using model design=∼Factor+Treatment and contrast=c(“Treatment”, “DEL”, “WT”). “Factor” represents the sample genotype (del1, del2 or WT), “Treatment” represents combined genotype, either DEL or WT. Regions with an FDR<0.1 were considered significantly DARs (see Supplementary Information for more details). DARs from different backgrounds in the same cell type were then merged by taking their overlaps to make the final cell-type-specific DARs. DARs target genes were identified as gene promoters (2000 bp upstream to 500 bp downstream of the transcription start site) that overlapped with cell-type-specific DARs.

One gene promoter might overlap with more than one DARs and the DARs contained in the same promoter region might be regulated in opposite directions.

#### Transcription factor footprint

TOBIAS (v0.12.10)^71^ was used to call TF footprint from ATAC-seq. First, the final alignments (i.e. BAM files) for samples from the same genotype within the same sequencing batch were merged. The correction for Tn5 shifting was achieved by calling the “ATACorrect” function of TOBIAS on each merged BAM file, within the corresponding merged peak set (i.e. per cell type per iPSC background) generated in the previous step (see “Transcription factor footprint” section in Methods). This generated a corrected BIGWIG file for ATAC-seq coverage, which was further scored by calling the “FootprintScores” function of TOBIAS per genotype per sequencing batch. The scored BIGWIG files were merged across sequencing batches using the USCS script “bigWigMerge” to generate footprint profiles per genotype per iPSC background. Then the “BINDetect” function was used to predict TF binding using the sequence motifs recorded in the JASPER database and compare the differential bindings between genotypes. For each TF’s motif, this generated two levels of changes between genotypes, namely: 1) locus level, which compares footprint difference between genotypes for each specific genomic locus that matches the TF motif; and 2) genome-wide level, which summarizes the global footprint difference between genotypes across all the loci of that TF. A differentially bound transcription factor (DTF) was defined as the genome-wide summarized changes of any motif in a single TF either 1) ranked top 5% of the largest -log10(pvalue); or 2) top and bottom 5% of differential binding score between genotypes. Finally, overlapped DTFs from the GM08830 and MGH2069 backgrounds were used as the final DTFs for *POGZ* heterozygous.

#### Direct and Indirect POGZ regulation

To generate the list of genes directly regulated by POGZ, POGZ-occupied loci in the human fetal cortex^22^ were used. Genes whose promoter overlapped with the POGZ-occupied loci were annotated as the direct POGZ targets in the human cortex. To generate the list of genes indirectly regulated by POGZ, each individual binding site belonging to a DTF was compared between genotypes. Those binding sites with substantial changes (predicted bound in at least one condition of DEL and WT, and the log2 fold change ranked within top or bottom 10%) of each DTF were retained. Any of these retained sites further overlapping with a gene promoter was considered as an indirect POGZ target.

### Multi-Electrode Array (MEA) analysis

48-well MEA plates (Axion Biosystems, M768-tMEA-48B) were prepared by coating wells with 50 µL 0.1% Polyethylenimine (PEI) (Millipore-Sigma, 408727) dissolved in Borate Buffer (Boston Bioproducts, BB-66-500) and sterile filtered with a 0.2μm filter prior to use. Coated wells were incubated at 37ºC for 1hr. Wells were then washed 6-10x with tissue culture-grade water (Boston Bioproducts, 7732-18-5) to remove residual PEI solution. Once excess water was removed, plates were stored at RT overnight to dry out completely. To prepare iN for electrophysiology readings we largely followed manufacturer protocol with a few modifications. Briefly, TRA1+ hiPSC lines underwent 1 post-thaw passage prior to further manipulation. All hiPSC lines for MEA underwent Ngn2 transduction as a single batch to eliminate viral transduction batch effect using the same transduction and selection steps described in “<u>Induction of</u> *<u>Ngn2</u>*<u>-neurons from hiPSC lines</u>” above. On Day 4 of iN differentiation, cells were detached using Accutase (STEMCELL, 07920), counted using a Countess Cell Counter (Thermo Fisher Scientific, A27977), and resuspended in NMM supplemented with BDNF, NT-3, Doxycycline and Laminin as outlined in manufacturer protocols (Axion Biosystems: Culturing Human hiPSC-derived Excitatory Neurons on Microelectrode Arrays: Maestro Pro MEA) in order to distribute 6x10^4^ cells per well in a volume of 10 µL directly to the center of each MEA plate well. Plated cells were left to adhere for 1hr at 37ºC prior to the addition of 200 µL NMM supplemented with BDNF, NT-3, and Doxycycline per well. 200 µL NMM supplemented with BDNF, NT-3, Doxycycline, and Ara-C was added on Day 5 for a total volume of 400 µL per well. Each well underwent a half media change (removal and addition of 200 µL) every 3 days with NMM supplemented with BDNF and NT-3 until Day 24. Doxycycline use was discontinued after Day 10 and Ara-C was only added on Day 6. Day 27 – Day 51 cells continued to undergo half media change every 3 days, but media was changed to BrainPhys (following manufacturer protocol: Axion Biosystems: Culturing Human hiPSC-derived Excitatory Neurons on Microelectrode Arrays: Maestro Pro MEA) to promote spiking activity. MEA plate readings were conducted every 3 days, 22-24 hours following each media change. Readings were conducted at 37ºC for 15min using Axion Biosystems Maestro Pro. Manufacturer set thresholds for spike and network burst calling were used. For data interpretation, only wells with at least 4 active electrodes were filtered. A total of 9 readings were performed from Day 24 to Day 51 and the reading with the highest number of clones with at least 4 active electrodes across all wells and genotypes was selected for statistical analysis (**Supplementary Figure 17**). Weighted mean firing rate was compared between genotypes using the Wilcoxon test.

### Incucyte analysis

The inner 60 wells of 96-well plates were prepared by coating wells with PLO/Laminin and iN were prepared for Incucyte recordings following the protocol described in “*<u>Ngn2-induced</u>*<u>neurons from hiPSC lines</u>” above, with volume adjustments. All hiPSC lines for Incucyte underwent Ngn2 transduction at the same batch performed for MEA readings to eliminate viral transduction batch effect. On Day 4 of iN differentiation, cells were detached using Accutase (STEMCELL, 07920), counted using a Countess Cell Counter (ThermoFisher, A27977), and resuspended in NMM supplemented with BDNF, NT-3 and Doxycycline in order to distribute 12x10^3^ cells in a volume of 75 µL per well. 75 µL NMM supplemented with BDNF, NT-3, Doxycycline, and Ara-C was added on Day 5 for a total volume of 400 µL per well. Each well underwent a half media change (removal and addition of 75 µL) every 3 days with NMM supplemented with BDNF and NT-3 until Day 27. Doxycycline use was discontinued after Day 10 and Ara-C was only added on Day 6. Stain-free Incucyte recordings were performed using a IncuCyte S3 with images acquisition every four hours, in three timepoints: Days 11-13, Days 18-20, and Days 25-27. Neurotrack analysis was performed in IncuCyte software (version 2020B) largely following default parameters, with a few modifications to increase sensitivity in detecting iN neurites and omit cellular debris. Neurotrack settings were “minimal cell width”=11, “minimal cell body cluster”=125, and “neurite sensitivity”=0.7. Neurite length and neurite branch points metrics were normalized by the area of cell body clusters in each well to eliminate variability due to artifactual cell counting differences. A total of 9 readings were performed from Day 11 to Day 27 and the reading with the highest overall neurite length across all wells and genotypes was selected for statistical analysis. Neurite length and neurite branch points were compared between genotypes using the Wilcoxon test.

## DECLARATIONS

## Acknowledgements

This research was supported by grants from the National Institutes of Health (NIH, R01MH123155, 5R01MH115957, R01HD096326, RM1MH132648, U01HG011755), the Simons Foundation Autism Research Initiative (SFARI, #573206 and #1009802). M.M.O. was supported by Autism Speaks Postdoctoral Fellowship (11815). R.Y. was supported by the Mass General Hospital Fund for Medical Discovery. P.M.B. was supported by the NIH (K08NS117891). D.G. was supported by the NIH (K99NS118109, R00NS118109-03).

## Authors’ contribution

Study design: M.M.O., Y.L., S.E., D.G., D.R., K.B., J.F.G., M.E.T. Experiments: M.M.O., Y.L., S.E., D.G., R.B., K.M., K.O.K, P.M.B., G.X., C.L., A.L., R.Y., M.S., D.L., B.C., C.E.F.E., D.J.C.T., D.R., K.B., J.F.G., M.E.T. Data interpretation: M.M.O., Y.L., S.E., D.G., D.R., K.B., J.F.G., M.E.T. Manuscript writing: M.M.O., Y.L., S.E., D.G., J.F.G., M.E.T. All authors have reviewed and accepted final version of the manuscript.

## Declaration of interests

M.E.T. receives research funding and/or reagents from Levo Therapeutics, Microsoft Inc, and Illumina Inc, is a paid consultant for BridgeBio, and is co-founder of First Genomic Insights, LLC. J.F.G. is a paid consultant to Harness Therapeutics and has consulted for Biogen, Inc., Pfizer, Inc., and Wave Biosciences, Inc. in the past five years.

## SUPPLEMENTAL MATERIAL

### Supplemental Information

- POGZ downregulation from RNA-seq data
- PCA and SV correction description in each cell type, justifying strategies for data integration in iN and in NSC
- WGCNA for POGZ models – modules description
- Differential accessibility regions (DARs) for *POGZ* cellular models
- TF expressed in NSC and iN and DTFs reproducibility across backgrounds
- Detection of DTF regulatory targets and their reproducibility across backgrounds DEG comparisons between *POGZ, MEF2C* and *SCN2A* heterozygous neuronal models

### Supplemental Figures

- Supplementary Figures 1-25

### Supplemental Tables

- Supplementary Tables 1-15

