## Supplemental Material for "CRISPR-engineered deletion of *POGZ* alters transcription factor binding at promoters of genes involved in synaptic signaling"

#### SUPPLEMENTAL INFORMATION

##### POGZ *downregulation from RNAseq data*

Using two different hiPSC backgrounds from healthy donors, we generated 23 independent CRISPR-edits, including heterozygous and compound heterozygous deletions of *POGZ* (**Supplementary Table 13**). Six hiPSC replicates per edit type and background, and six matched controls, were differentiated into NSC and iN for transcriptional profiling, totalizing 132 RNAseq libraries (**Supplementary Table 13**). POGZ copy number as well as downregulation at protein and transcriptional levels were validated in hiPSCs and hiPSC-derived neural lineages (**Supplementary Figure 1** and **2**). Evaluation of cell type-specific markers confirmed the successful hiPSC differentiation into the targeted neuronal models (**Supplementary Figure 18**).

##### *PCA and SV correction description in each cell type, justifying strategies for data integration in iN and in NSC*

To investigate the variation between samples due to known and unknown sources, principal component analysis (PCA) was performed on all the RNAseq libraries. PCA revealed the largest variation between samples was caused by background in iN samples, as the first principal component (PC1) accounting for 72% variation showed two distinct clusters by background, whereas the second principal component (PC2) accounting for 9% variation seems to separate batch effects within each background (**Supplementary Figure 19**). This result urged us to perform differential expression analysis within each background, combining samples from different batches. To this end, surrogate variables estimated by R/Bioconductor package, SVA 36 was used in differential expression models to correct for batch effects and unknown sources of variation. PCA within each background in iN based on SVA-corrected counts revealed that PC1 separated samples by genotype, accounting for 16-19% variation (**Supplementary Figure 19**). iN DEGs comprised genes that were consistently differentially expressed at FDR < 0.1 in both backgrounds. We followed a different approach to identify DEGs in NSC, since PCA on NSC samples revealed that MGH2069 del2 batch was very distant from other NSC batches (**Supplementary Figure 20**). Moreover, NSC samples were not separated by genotype after SVA correction of combined samples within each background (**Supplementary Figure 20**). To identify DEGs in each background, we performed meta-analysis following weighted-Stouffer's z-score method for which we first generated a z-score for each gene that passed the expression threshold in each batch of a particular background. Gene-level meta z-scores were computed from these z-scores for each background and were further converted to p-values following normal distribution. Adjusted p-values (FDR) were calculated separately for each background, following Benjamini-Hochberg procedure. Genes dysregulated in the same direction in each batch for a particular background at a significance level of FDR < 0.1 were defined as DEGs. The consensus DEGs for NSC were defined following the same criteria described above for iN.

PCA of two batches consisting of 17 NSC samples (6 wild-type, 6 del1, 5 compound heterozygous (del1-del2)) and 18 iN samples (6 wild-type, 6 del1, 6 del1-del2) from GM08330 background revealed that del1-del2, del1 and wild-type samples were distinctly clustered in the first and second PCs accounting for 70% and 17% variance in NSC and 75% and 15% variance in iN (**Supplementary Figure 21**). Since PCA showed a clear separation of both NSC and iN samples from each genotype, SVA 36 was not used in differential expression analysis of del1-del2 vs wild-type and del1 vs wild-type comparisons performed by DESeq2 78.

##### *WGCNA for* POGZ *models – modules description*

All batches in iN were used for WGCNA analysis. For NSC, we removed MGHdel2 batch (6 del2 and 6 wt in MGH2069 background), because of their distance from other batches in PCA plots.

Co-expression analysis of iN *POGZ* Het and WT samples identified 37 modules with sizes varying between 54 and 1,620. *POGZ* was assigned to iN module “red”. Co-expression analysis of NSC *POGZ* Het and WT identified 18 modules with sizes varying between 65 and 1,783. *POGZ* was assigned to NSC module “blue”. Co-expression analysis of iN *POGZ* Het, *SCN2A* Het and WT samples identified 29 modules with sizes varying between 51 and 1,075. *SCN2A* was assigned to “turquoise” module, whereas *POGZ* was not assigned to any module after filtering based on module membership p-values. To identify the modules of interest in co-expression analysis of iN *POGZ* and *SCN2A* samples, module eigengenes were fitted against the following four contrasts using linear regression; *POGZ* Het vs *SCN2A* Het + WT, *SCN2A* Het vs *POGZ* Het + WT, *POGZ* Het + *SCN2A* Het vs WT (concordant), a contrast named “discordant” in which *POGZ* Het, *SCN2A* Het and WT samples were assigned to (1,-1,0). Additionally, module eigengene values for *POGZ* Het vs WT and *SCN2A* Het vs WT comparisons were tested using two-tailed t-test for each of 29 modules. Overall, 6x29=174 tests were performed. Following Bonferroni multiple-test correction procedure, adjusted p-value cut off was set to 0.05/174=0.000287. In the first pass, 15 modules passed this threshold in at least one of six tests. Next, modules were tested for enrichment of up-regulated and down-regulated DEGs ay FDR<0.1 from *POGZ* Het vs WT (only using samples from GM08330 as we used for *POGZ+SCN2A* co-expression analysis) and *SCN2A* Het vs WT comparisons using one-tailed Fisher’s exact test. In doing so, each module was tested for four distinct DEG lists. Therefore, in total 4x29=116 tests were performed. Bonferroni adjusted p-value was set to 0.05/116=0.000431. Fourteen out of 15 modules selected in the first pass were enriched for at least one DEG list, which are depicted in **Figure 2C**.

#### Differential accessibility regions (DARs) for POGZ cellular models

We first generated four ATAC-peak profiles according to cell types and hiPSC backgrounds, namely, iN-GM08330, iN-MGH2069, NSC-GM08330, and NSC-MGH2069. Within each profile, DARs of “del1-vs-WT” and “del2-vs-WT” were performed independently. In iN, the DARs derived from “del1-vs-WT” and “del2-vs-WT” are highly correlated in both backgrounds (data not shown), so we merged “del1” and “del2” into a simple heterozygous deletion condition. This resulted in 293 and 220 genome-wide significant DARs (FDR < 0.1) for iN-GM08330 and iN-MGH2069, respectively. This FDR cutoff represented the top 0.18% peak changes. In NSC, surprisingly, “del2-vs-WT” in both backgrounds did not lead to any DAR at the FDR level. Given that the aim was to study the POGZ’s epigenetic regulation, we considered “del2” unsuitable for the downstream analyses and removed it. “del1-vs-WT” in NSC-MGH2069 displayed 255 genome-wide significant DARs at the FDR < 0.1, representing top 0.17% peak changes. “del1-vs-WT” in NSC-GM08330, however, had 61,214 genome-wide significant DARs (FDR < 0.1). A further investigation of FDR distribution indicated it was inflated (**Supplementary Figure 22A**). To control this inflation, we only considered the top 0.18% peak changes as its DAR, a level comparable to DARs in other conditions. The final genome-wide DARs and their overlaps with promoters were provided in **Supplementary Table 8**. When we investigated DARs’ enrichment in promoters of DEGs we noticed that POGZ-target DEGs did not show a stronger association with DAR than non-POGZ-target DEGs (**Supplementary Figure 22B**).

Consistent with the extensive transcriptomic effects (**Figure 4A**), *POGZ* compound heterozygosity caused major changes in global promoter accessibility, which was increased in iN but decreased in NSC; thus, in opposite directions in the two cell types (**Supplementary Figure 23**). Consequently, in the compound heterozygous state, iN displayed far more DARs than NSC (N=158,027 versus 917, **Supplementary Table 8**). Genes whose promoters were affected by DARs were highly enriched for DEGs, with gene expression and DNA accessibility generally showing a consistent direction of effect (i.e., the promoters of upregulated DEGs showed higher DNA accessibility and vice versa, **Figure 23A-D**). However, in compound heterozygous lines, both upregulated and downregulated POGZ targets were enriched among genes regulated by promoters with increased DNA accessibility in iN (**Supplementary Figure 23C and D**). The upregulated POGZ targets with increased accessibility were enriched for genes involved in processes related to synapse, extracellular matrix, immune pathways, RNA processing, cell cycle, transcription and translation-related processes, whereas those downregulated implicated a subset of these processes (e.g., RNA processing) (**Supplementary Figure 23D**). Taken together, these experiments indicate that *POGZ* LoF can alter chromatin accessibility on promoters of genes directly regulated by POGZ. However, such changes in DNA accessibility over POGZ targets did not necessarily impact gene expression modulation, suggesting that POGZ might act in conjunction with other transcription regulators.

#### TF expressed in NSC and iN and DTFs reproducibility across backgrounds

In both cell types, differentially “footprinted” transcription factors (DTFs) were significantly overlapped between background lines from different donors (i.e. MGH2069 and GM08330) (**Supplementary Figure 24**), assuring the reproducibility of our experiments and analysis pipelines. We considered as biologically relevant only those factors which showed significant altered footprinting with concordant direction of effect between backgrounds (see methods).

#### Detection of DTF regulatory targets and their reproducibility across backgrounds

After detecting DTF between genotypes (see Methods for details), in each iPSC background (i.e. MGH2069 and GM08330) we defined DTF regulatory targets as the genes immediately downstream to those promoters containing the DTF sites. By doing so, we found that the DTF regulatory targets between the two hiPSC backgrounds were significantly shared between the two backgrounds. In iN, there were 3,676 (~55%) shared targets from 6,627 targets in MGH2069 and 6,225 targets in GM08330, respectively ($p$=9.97e-188, hypergeometric test). Similarly, in NSC, we found 1628 (~35%) shared targets from 4,064 targets in MGH2069 and 4,666 targets in GM08330, respectively (p=1.99e-71, hypergeometric test). These results suggested the DTF regulatory targets we identified are consensus to genotypes and not biased to confounding factors such as batch or background effects.

#### DEG comparisons between POGZ, MEF2C and SCN2A heterozygous neuronal models

We compared DEG across *POGZ, MEF2C* and *SCN2A* heterozygous iN models with gene deletions. representing two chromatin-related genes, and *SCN2A*, representing genes involved in synaptic function. The subset of DEGs shared across all three ASD/NDD gene models (filtered at FDR < 0.3) was enriched for signaling pathways, including cytokine and Eph signaling, LoF constraint genes and NDD-risk genes (**Figure 25**).

### SUPPLEMENTAL FIGURES

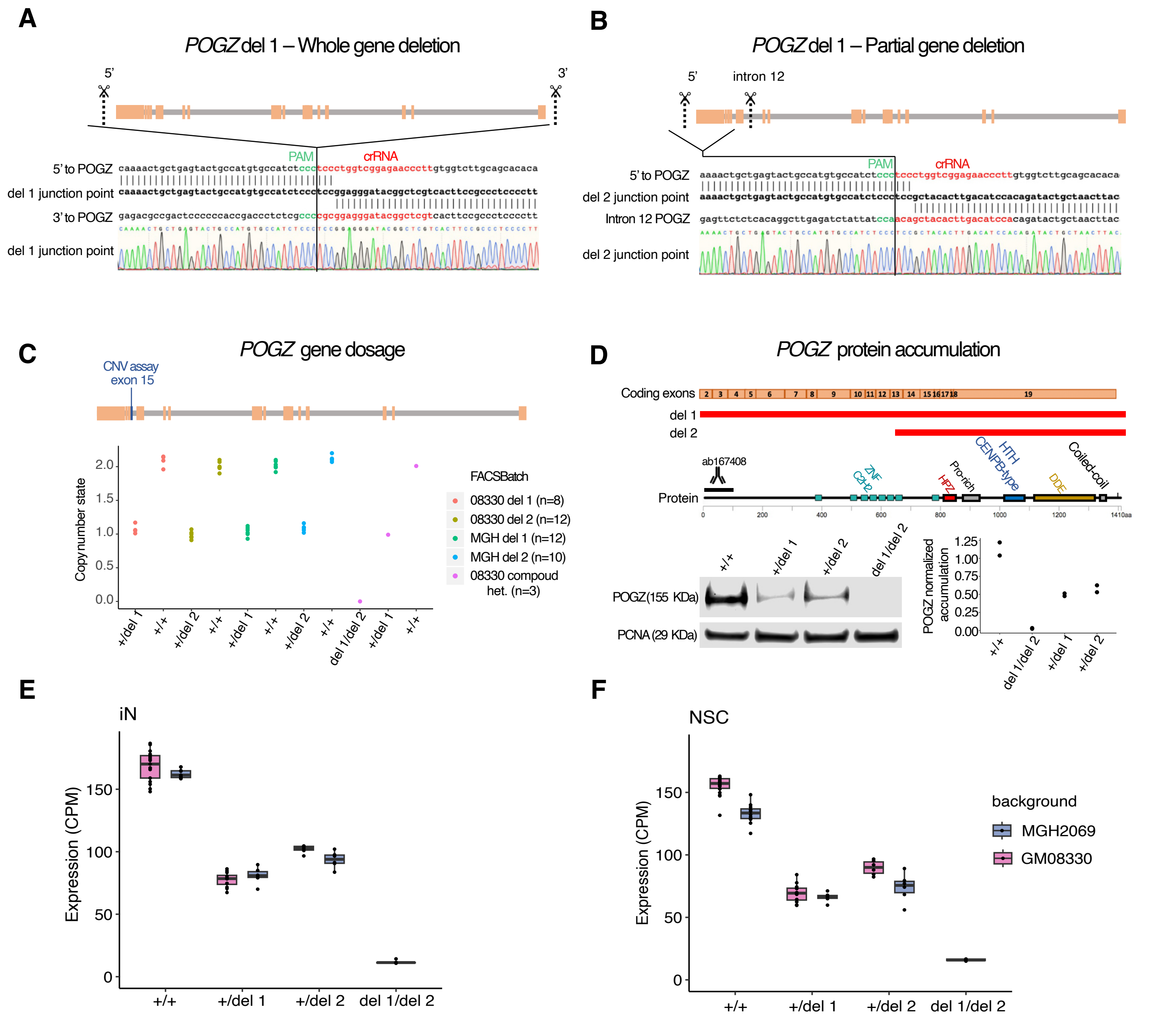


***Supplementary Figure 1. CRISPR editing design to generate POGZ deletions and validation of POGZ loss of function in resultant cell lines.***

**(A)** and **(B).** Representations of gRNA targets localization (5' to POGZ, intron 12 and 3' to POGZ) and combination of gRNAs used to generate the whole gene deletion (Del 1, panel A) and the partial gene deletion (Del 2, panel B). Sanger sequencing of the deletions' junction points aligned to wild-type gRNA target regions, showing examples of Sanger traces from clones with no insertions or indels at cut sites. PAM and crRNA sequences are indicated in green and red, respectively. **(C)** Determination of *POGZ* gene dosage in heterozygous (+/del1 and +/del2), compound heterozygous (del1/del2) and wild-type control (+/+) hiPSC lines using TaqMan copy number probes on a ddPCR assay. The Taqman copy-number variant (CNV) assay used in this experiment targeted *POGZ* at exon 15. CRISPR transfection/ FACS batches are labeled by color code. **(D)** Representation of POGZ coding exons and the corresponding encoded protein domains, indicating the protein segments affected by each deletion (Del 1 and Del 2). POGZ protein expression in wild type control (+/+), heterozygous (+/del1 and +/del2), and compound heterozygous (del1/del2) NSC lines by western blot analysis using a POGZ primary antibody with a N-terminal epitope (ab171934). PCNA protein expression was used to normalize POGZ signal. *POGZ* isoform NM_015100 used as reference. Western blot experiments were performed twice. **(E)** and **(F).** mRNA expression level of *POGZ* transcripts in iN (E) and NSC (F) lines on MGH (blue) and GM (pink) backgrounds with different edit types and zygosities. WT: wild-type, del1: heterozygous full *POGZ* gene deletion, del2: heterozygous partial *POGZ* gene deletion, del1_del2: compound heterozygous.

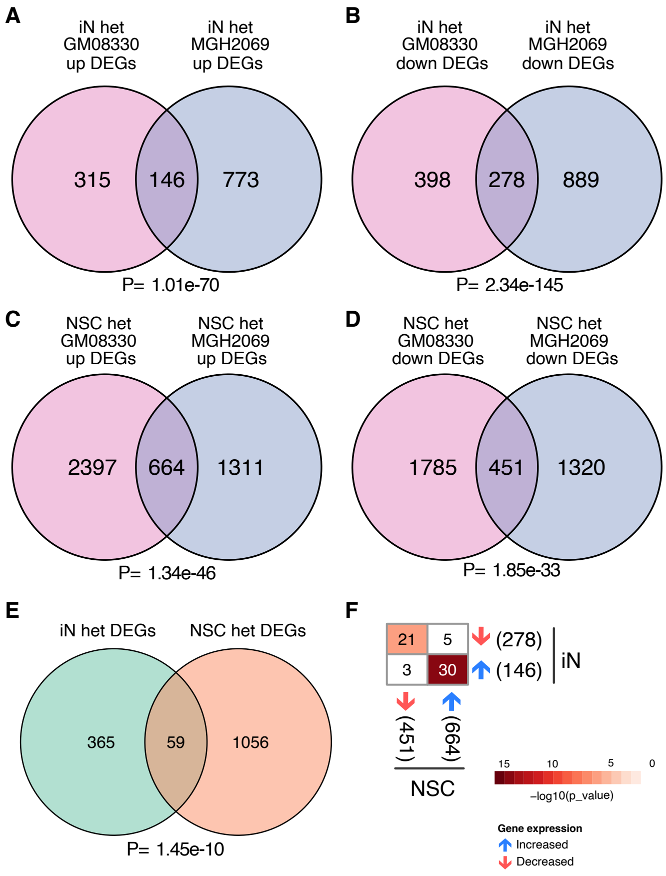


***Supplementary Figure 2. Overlap between DEGs in different background lines, cell types and zygosities***

**(A)** to **(D)** Overlap of DEGs in concordant direction of effect between GM (pink) and MGH (blue) background lines in iN (panel A and B) and NSC (panel C and D).
**(E)** Overlap between DEGs in iN (green) and NSC (orange) het models.

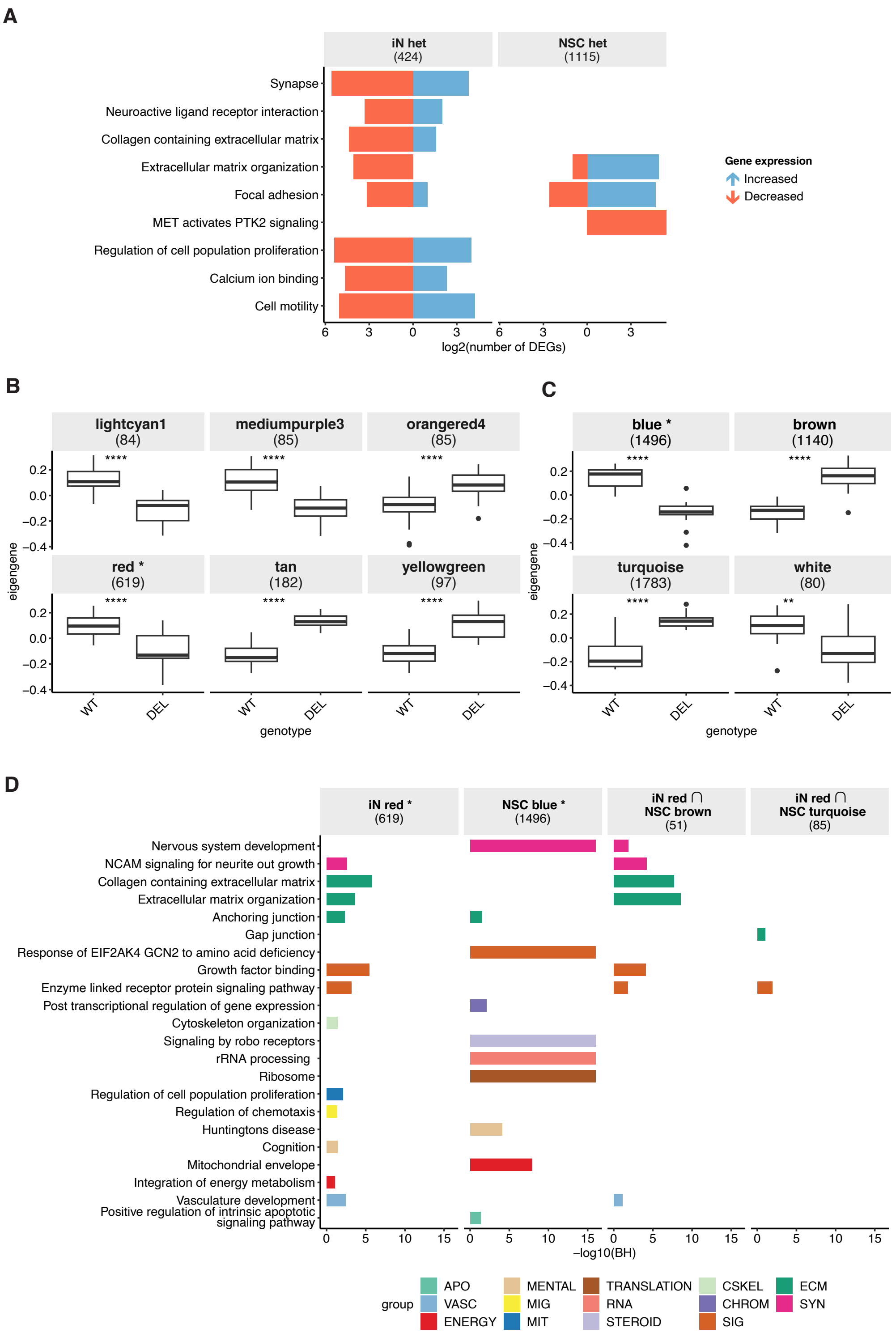


***Supplementary Figure 3. Co-expression modules associated with POGZ genotype in heterozygous hiPSC-derived iN and NSC***

**(A)** Number of upregulated (blue) and downregulated (red) genes that are part of the enriched pathways among DEGs in het iN and NSC lines. **(B)** and **(C)** Eigengenes expression levels for each module associated with POGZ genotype in iN (panel A) and in NSC (panel B). Asterisks indicate the module containing POGZ and numbers in parentheses indicate the total number of genes in the module. **(D)** Statistical significance for enrichment of pathways among genes in co-expression modules containing POGZ (asterisk): iN red and NSC blue, and among genes shared between iN red and either of the NSC POGZ genotype-correlated modules (brown and turquoise). Intersection of two gene lists is represented by "∩". Numbers in parentheses indicate the total number of genes in a given gene list. Colors represent pathway categorization according to the nature of the functional categories related to the enriched term. *Groups of functional categories: APO: apoptosis, VASC: vascular system, ENERGY: energy metabolism, MENTAL: mental and behavioral traits, MIG: cell migration, MIT: mitosis and cell proliferation, TRANSLATION: translation, RNA: RNA metabolism, STEROID: steroid metabolism, CSKEL: cytoskeleton, CHROM: chromatin structure and regulation, SIG: intracellular signaling pathways, ECM: extracellular matrix, SYN: synapse*. BH: Benjamini-Hochberg adjusted p-values.

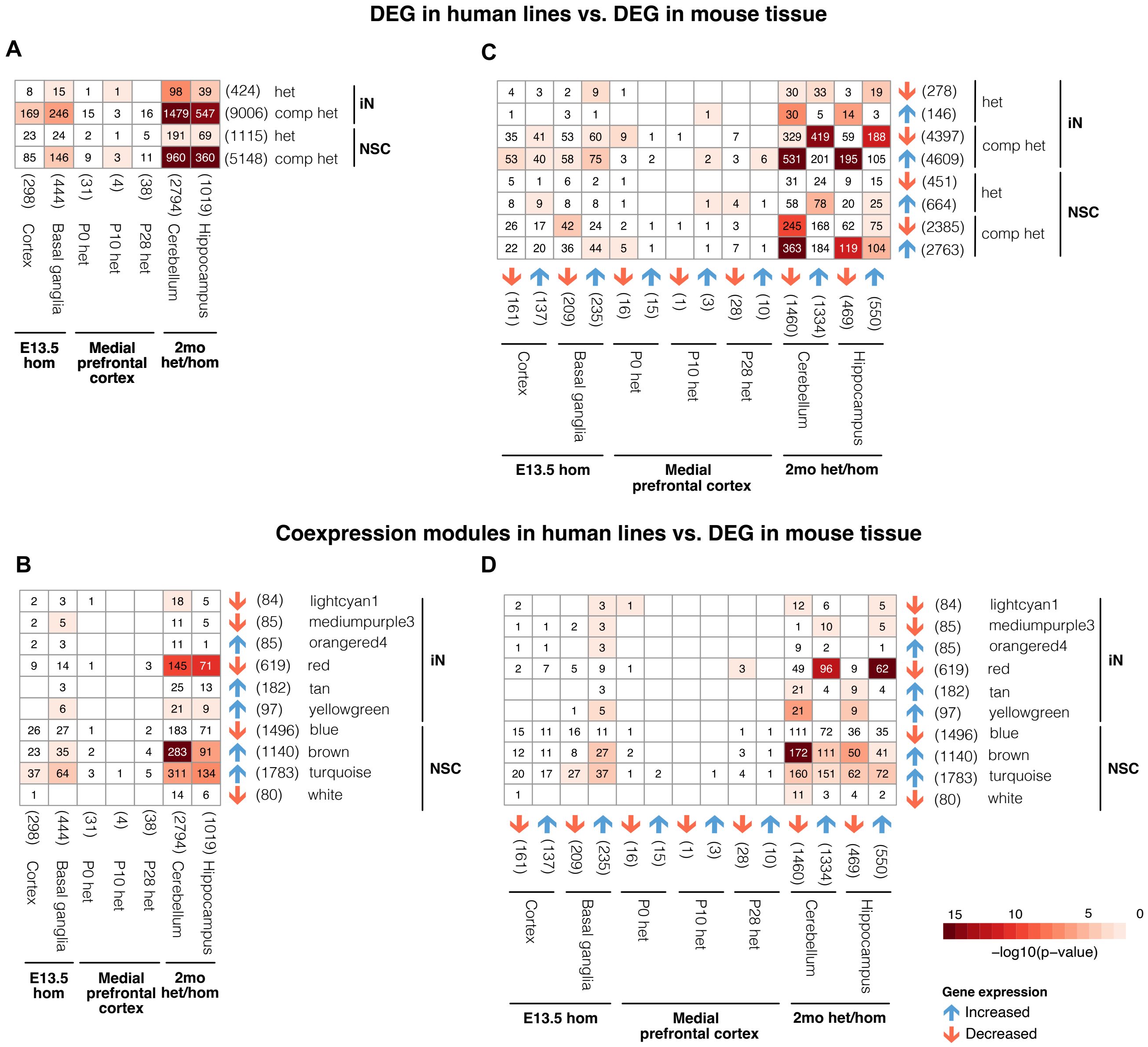


***Supplementary Figure 4. Overlap between transcriptional profiles from POGZ LoF models in hiPSC-derived neural lineages and mouse pre and postnatal brain tissues***

**(A)** and **(B)** Heatmap indicating statistical significance for overlap between human iN or NSC DEGs from heterozygous lines with (het: heterozygous, up: DEG with increased gene expression in edited cell lines, down: DEG with decreased gene expression in edited cell lines) and mouse DEGs from pre and postnatal brain tissues. Panel A considers overall DEG lists, and panel B considers DEG' direction of effect. **(C)** and **(D).** Heatmap indicating statistical significance for overlap between co-expression modules in human heterozygous iN or NSC and mouse DEGs from pre and postnatal brain tissues. Panel C considers overall DEG lists, and panel D considers DEG direction of effect. Numbers in parentheses indicate the total number of genes in a given gene list. Asterisks indicate *POGZ*-containing modules.

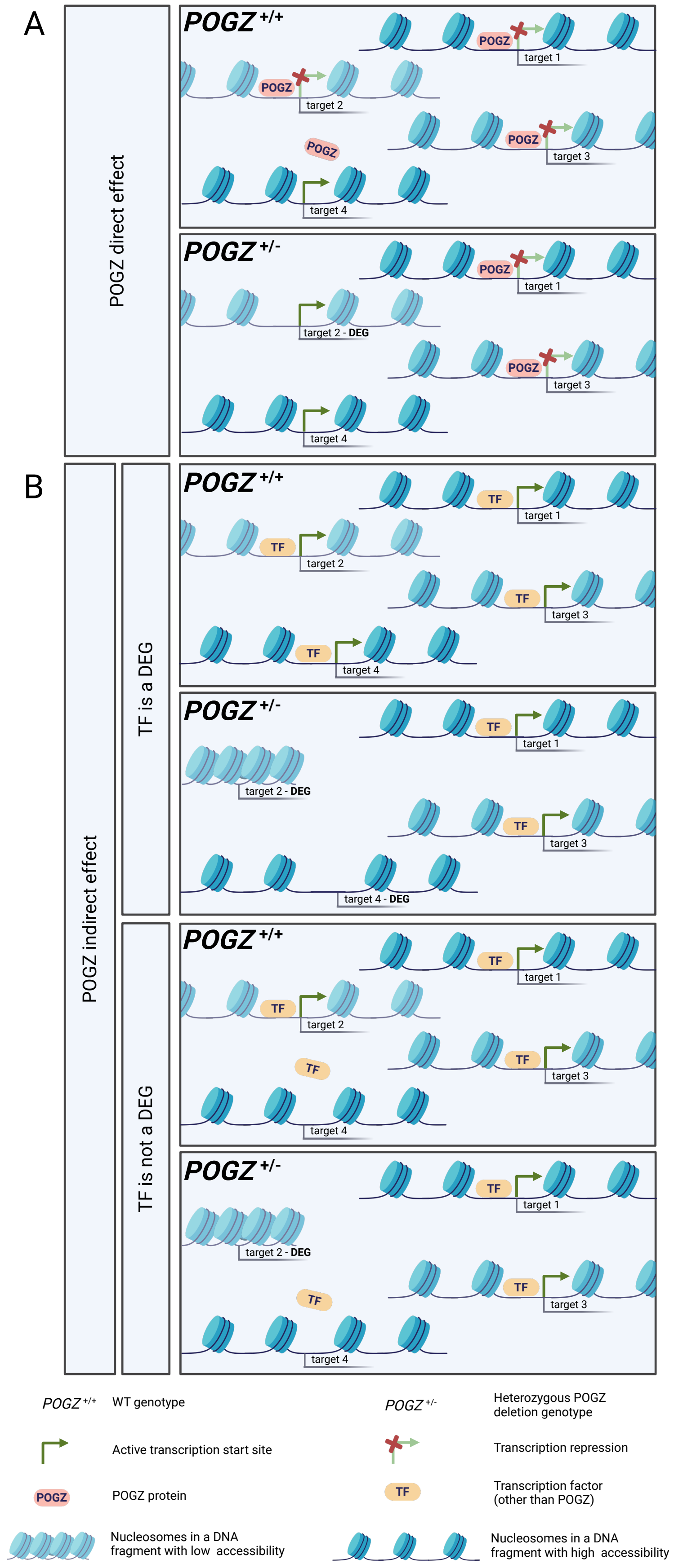


***Supplementary Figure 5. Scheme with definitions of direct and indirect effects of POGZ heterozygous loss of function over genes expression regulation***

**(A)** The direct effect is observed in differential expressed genes (DEGs) that are a POGZ target, such as "target 2", a POGZ regulatory target that is a DEG in *POGZ*$^{+/-}$ lines. **(B)** The indirect effect is observed in two situations. The first situation (upper panel in B), *POGZ*$^{+/-}$ DEGs are targets of a differentially expressed TF - other than POGZ. Target 2 and target 4 represent this situation. The second situation (bottom panel in B), *POGZ*$^{+/-}$ DEGs are targets of a TF - other than POGZ - that is not differentially expressed but presents with differential footprinting at its DNA binding sites. Target 2 represents this situation. Only the POGZ function as transcription repressor is being considered. DEG: differentially expressed genes, represented by arrows, TF is not a DEG: Shows a transcription factor that is not a differentially expressed gene. "TF is a DEG": Shows a transcription factor that is a differentially expressed gene. POGZ$^{+/+}$: wild-type POGZ genotype. POGZ$^{+/-}$: heterozygous POGZ deletion. Target: transcription start site of a specific regulatory target gene. *Target – DEG: transcription start site of a specific regulatory target gene that is differentially expressed. Green arrows: gene expression. Red cross: repression of gene expression.*

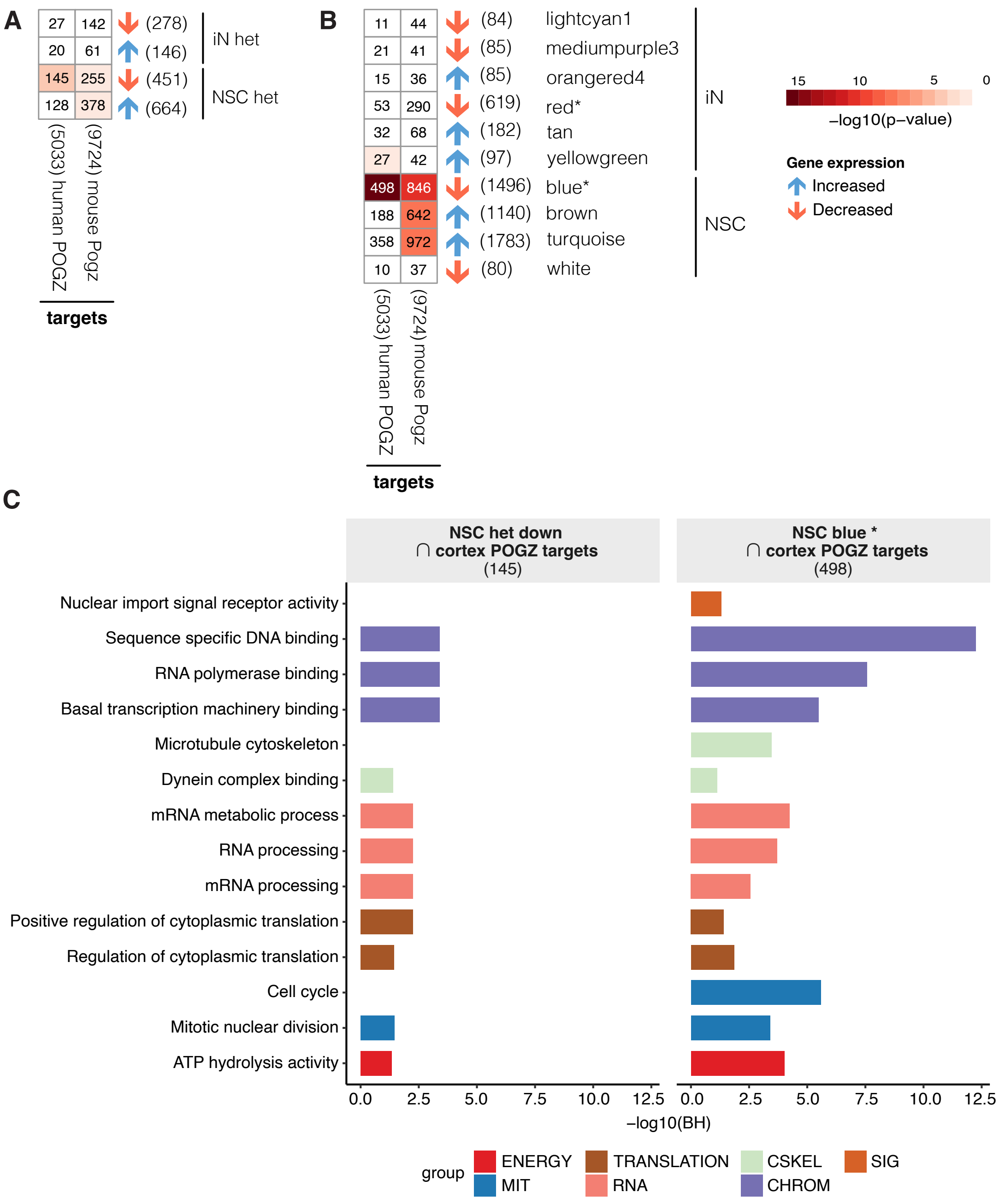


***Supplementary Figure 6. Overlap between transcriptional profiles and POGZ targets demonstrating POGZ direct effects on gene expression regulation***
(A) Heatmaps indicating statistical significance for overlap of iN or NSC up/downregulated DEGs from heterozygous lines and POGZ direct regulatory targets in human fetal cortex or mouse embryonic stem cells. (B) Heatmaps indicating statistical significance for overlap of iN or NSC co-expression modules associated with genotype in heterozygous lines and POGZ direct regulatory targets in human fetal cortex or in mouse embryonic stem cells. (C) Statistical significance for enrichment of pathways among human fetal cortex POGZ direct target genes that are also downregulated DEGs in heterozygous NSC (left) or are found in the NSC co-expression module containing POGZ (right). Intersection of two gene lists is represented by "∩". Numbers in parentheses indicate the total number of genes in a given gene list. Asterisk indicates POGZ containing module. Colors represent pathway categorization according to the nature of the functional categories related to the enriched term. *Groups of functional categories: ENERGY: energy metabolism, MIT: mitosis and cell proliferation, TRANSLATION: translation, RNA: RNA metabolism, CSKEL: cytoskeleton, CHROM: chromatin structure and regulation, SIG: intracellular signaling pathways.* BH: Benjamini-Hochberg adjusted p-values.

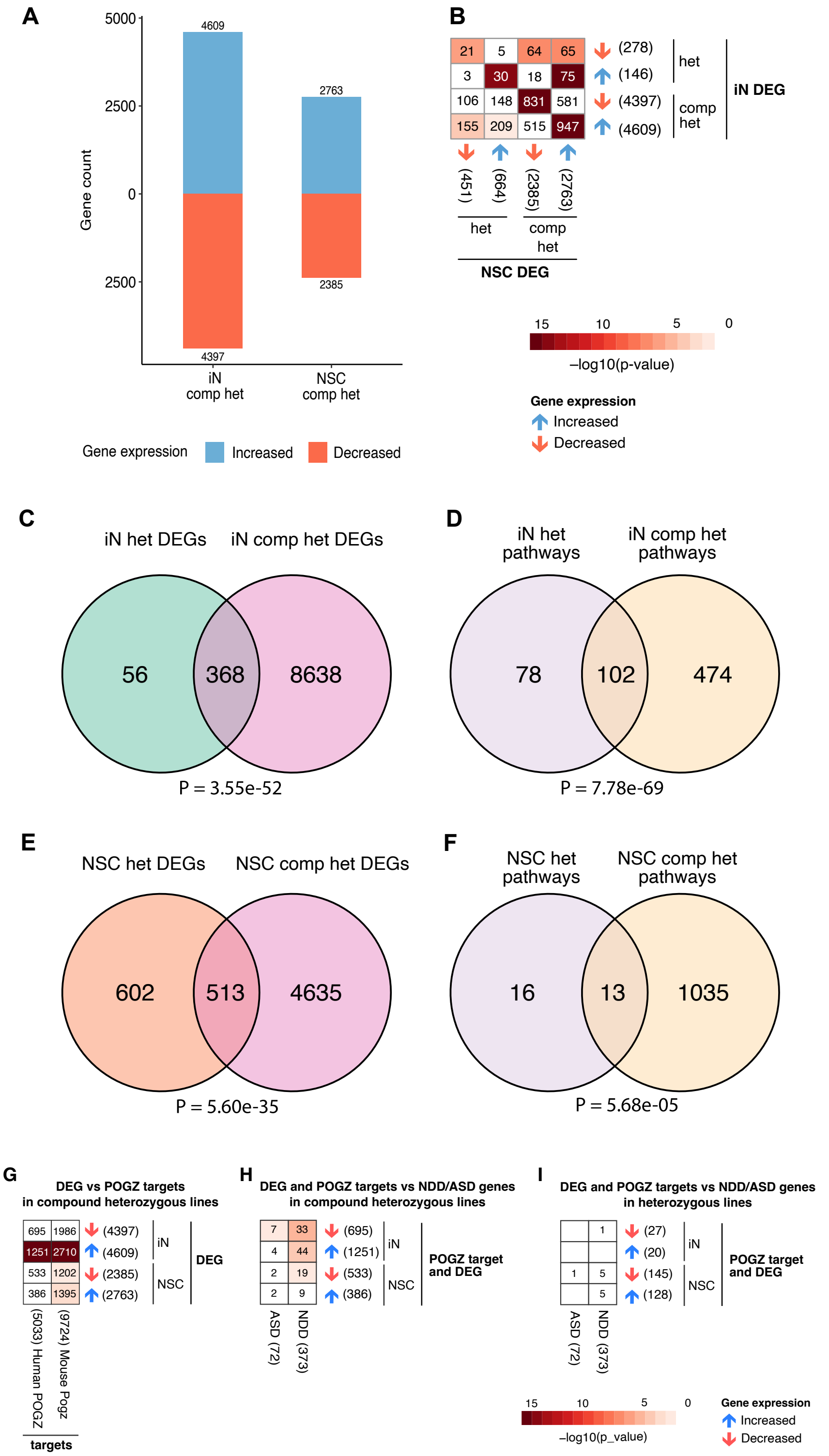


***Supplementary Figure 7: Transcriptional profiles of NSC and iN lines with compound heterozygous POGZ deletion***

**(A)** Number of upregulated (blue) and downregulated (red) genes in iN and NSC compound heterozygous (comp het) models. B. Heatmap indicating statistical significance for overlap between iN or NSC up/downregulated DEGs from lines with different zygosities (het: heterozygous, comp het: compound heterozygous, up: DEG with increased gene expression in edited cell lines, down: DEG with decreased gene expression in edited cell lines). Total genes indicated in brackets. C. Overlap between iN DEGs in heterozygous (het) models (green) and compound heterozygous (comp het) models (pink). D. Overlap between iN DEGs' enriched pathways in het models (purple) and comp het models (yellow). E. Overlap between NSC DEGs in het models (orange) and comp het models (pink). F Overlap between NSC DEGs' enriched pathways in het models (purple) and comp het models (yellow). G. Heatmaps indicating statistical significance for overlap of iN or NSC up/downregulated DEGs from compound heterozygous lines and POGZ direct regulatory targets in human fetal cortex or mouse embryonic stem cells. H and I. Heatmap indicating statistical significance for overlap between iN or NSC up/downregulated DEGs which are POGZ direct targets in human fetal cortex and genes associated with ASD and NDD in compound heterozygous (H) and in heterozygous (I) lines.

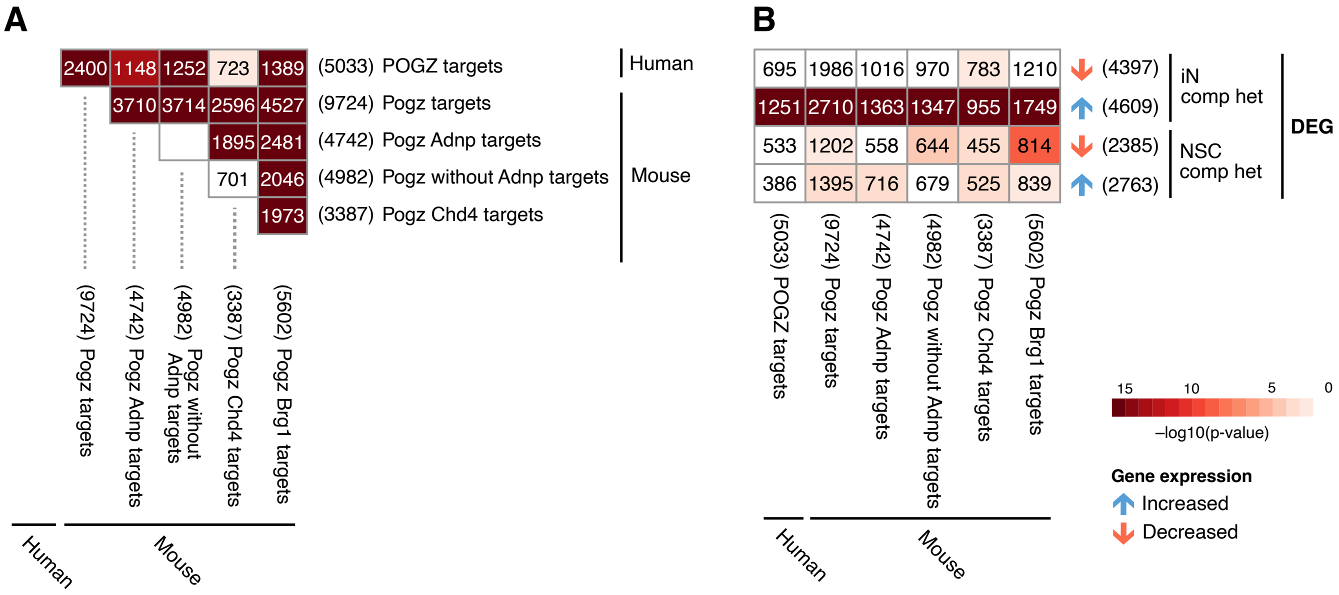


***Supplementary Figure 8. Effect of* POGZ *protein interactors on gene expression modulation in hiPSC-derived iN and NSC***

**(A)** Heatmap indicating statistical significance for overlap between POGZ direct targets identified in human fetal cortex and different subsets of Pogz direct targets identified in mouse embryonic stem cells. Subsets of Pogz direct targets in mouse cell lines are defined based on the Pogz protein interactors, as Adnp, Chd4 and Brg1. Only POGZ targets which are expressed in iN and NSC were considered. **(B)** Heatmap indicating statistical significance for overlap between iN or NSC up/downregulated DEGs from compound heterozygous lines (comp het: compound heterozygous, up: DEG with increased gene expression in edited cell lines, down: DEG with decreased gene expression in edited cell lines) and different subsets of POGZ/Pogz direct regulatory targets. Numbers in parentheses indicate total number of genes in a given gene list.

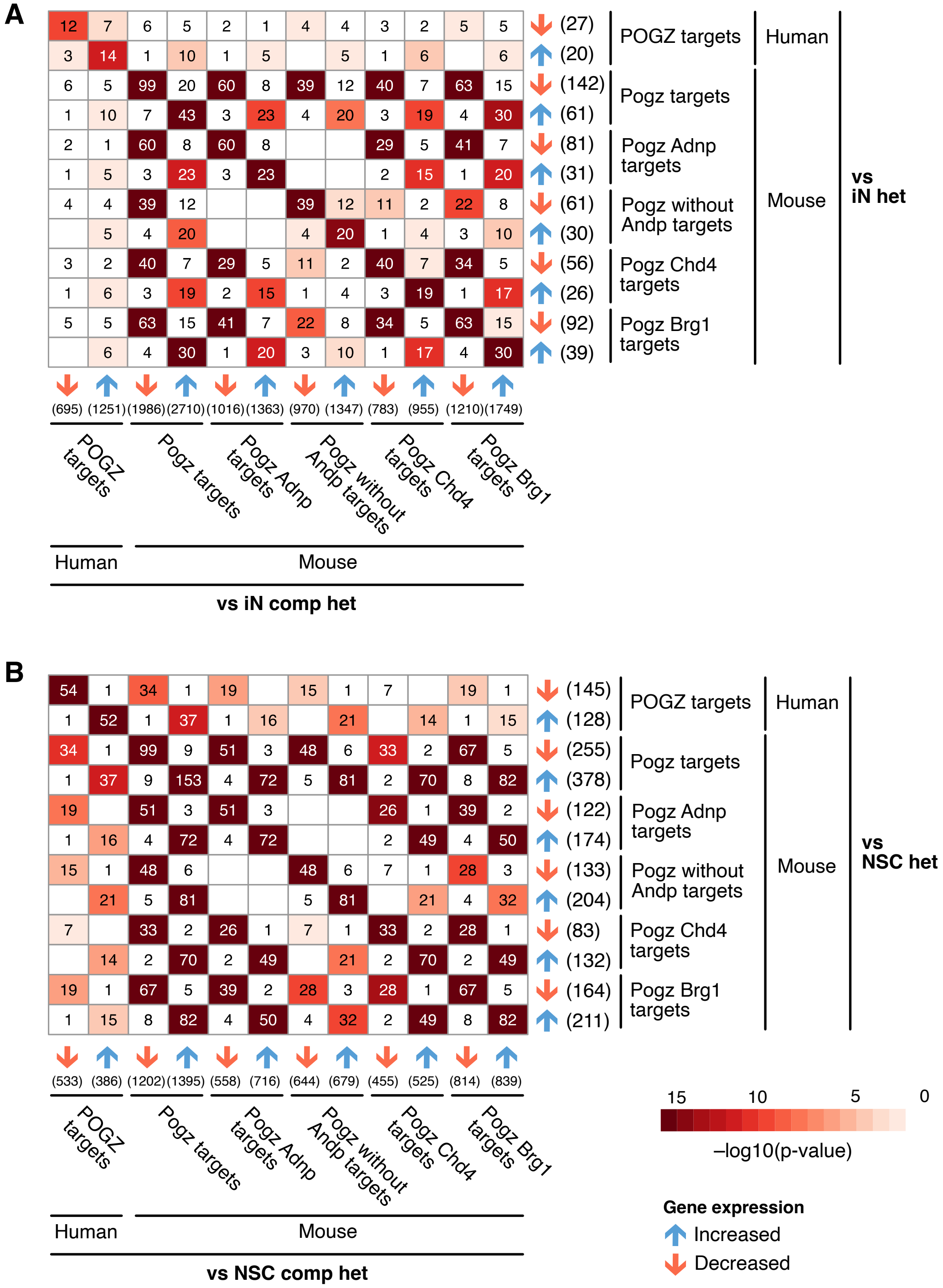


***Supplementary Figure 9. Effect of zygosity on gene expression modulation of POGZ direct targets***

**(A)** and **(B)**. Heatmap indicating statistical significance for overlap with POGZ direct targets which are up/downregulated DEGs in iN (panel A) and NSC (panel B) heterozygous (het) and compound heterozygous (comp het) lines. Different subsets of Pogz targets in mouse embryonic stem cells are determined based on Pogz protein interactors, as Adnp, Chd4 and Brg1. Numbers in parentheses indicate the total number of genes in a given gene list. *up: DEG with increased gene expression in edited cell lines, down: DEG with decreased gene expression in edited cell lines.*

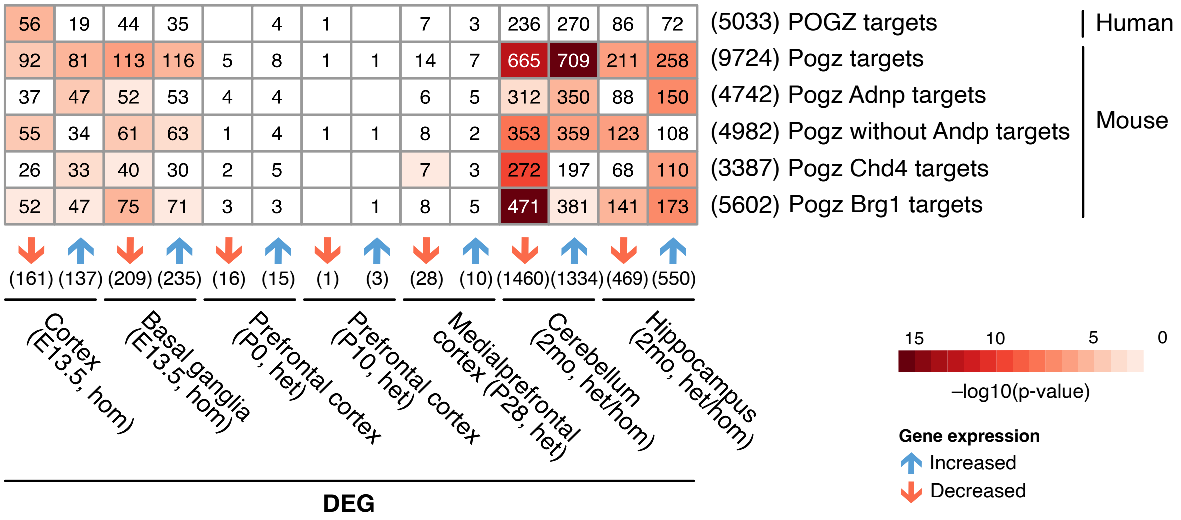


***Supplementary Figure 10. Effect of POGZ protein interactors on gene expression modulation in mouse models***

Heatmap indicating statistical significance for overlap between up/downregulated DEGs from mouse pre or postnatal brain tissues and different subsets of POGZ/Pogz direct regulatory targets. Subsets of Pogz targets in mouse embryonic stem cells are determined based on Pogz protein interactors, as Adnp, Chd4 and Brg1. Numbers in parentheses indicate the total number of genes in a given gene list.

### DAR mapped to gene promoters (DAR genes) vs. DEGs

| iN GM80330 (p= 2.823e-09) | DAR genes | Not DAR genes |
|---|---|---|
| DEGs | 13 | 1124 |
| Not DEGs | 13 | 15563 |

| iN MGH2369 (p=5.214e-13) | DAR genes | Not DAR genes |
|---|---|---|
| DEGs | 21 | 2065 |
| Not DEGs | 9 | 14484 |

| NSC G80330 (p=0.04108) | DAR genes | Not DAR genes |
|---|---|---|
| DEGs | 17 | 5280 |
| Not DEGs | 18 | 10696 |

| NSC MGH2369 (p=2.394e-06) | DAR genes | Not DAR genes |
|---|---|---|
| DEGs | 30 | 3716 |
| Not DEGs | 25 | 11271 |

| iN - backgrounds combined (p= 9.114e-10) | DAR genes | Not DAR genes |
|---|---|---|
| DEGs | 7 | 417 |
| Not DEGs | 3 | 15739 |

| NSC - backgrounds combined (p= 0.007139) | DAR genes | Not DAR genes |
|---|---|---|
| DEGs | 4 | 1111 |
| Not DEGs | 7 | 13570 |

***Supplementary Figure 11. Integration between differential expression and accessibility analysis with contingency tables for enrichment of differentially accessible regions (DAR) on promoter of differentially expressed genes (DEG)***

In the columns, genes were categorized as with or without DAR mapped to promoters ("DAR genes" and "Not DAR genes", respectively). In the rows, genes were categorized as DEG or not DEG. In each cell type, analysis was performed for each background separately (MGH2069 and GM08330) and combining both backgrounds.

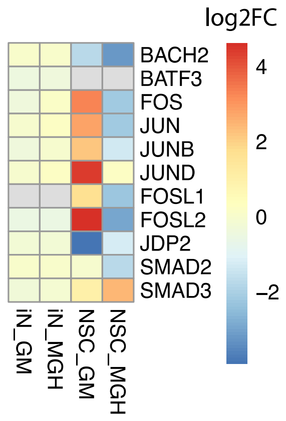


***Supplementary Figure 12. Integration of differential expression and footprinting analysis to identify indirect regulatory mechanisms***

Heatmap indicating gene expression level fold change of genes which encode DTF members in heterozygous iN and NSC from GM08330 and MGH2069 background lines.

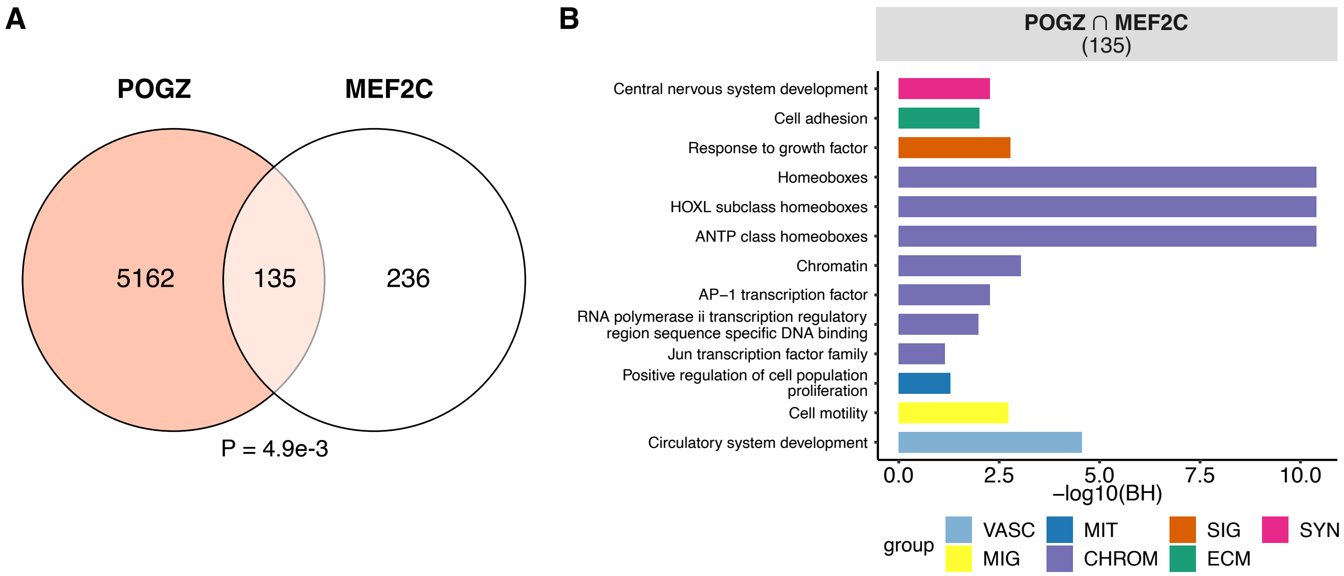


***Supplementary Figure 13. Overlap of transcriptional profiles derived from isogenic heterozygous NSC with POGZ or MEF2C gene deletions***

**(A)** Overlap between DEGs (FDR<0.1) from heterozygous NSC with POGZ or MEF2C gene deletions. **(B)** Statistical significance for enrichment of pathways among DEG overlaps between heterozygous NSC with POGZ or MEF2C gene deletions. Colors represent pathway categorization according to the nature of the biological process related to the enriched term. Numbers in parentheses indicate the total number of genes in a given gene list. Groups of biological processes: *IMMUNE: immune system and inflammation, CALCIUM: calcium signaling, SIG: intracellular signaling pathways, SYN: synapse.* BH: Benjamini-Hochberg adjusted p-values.

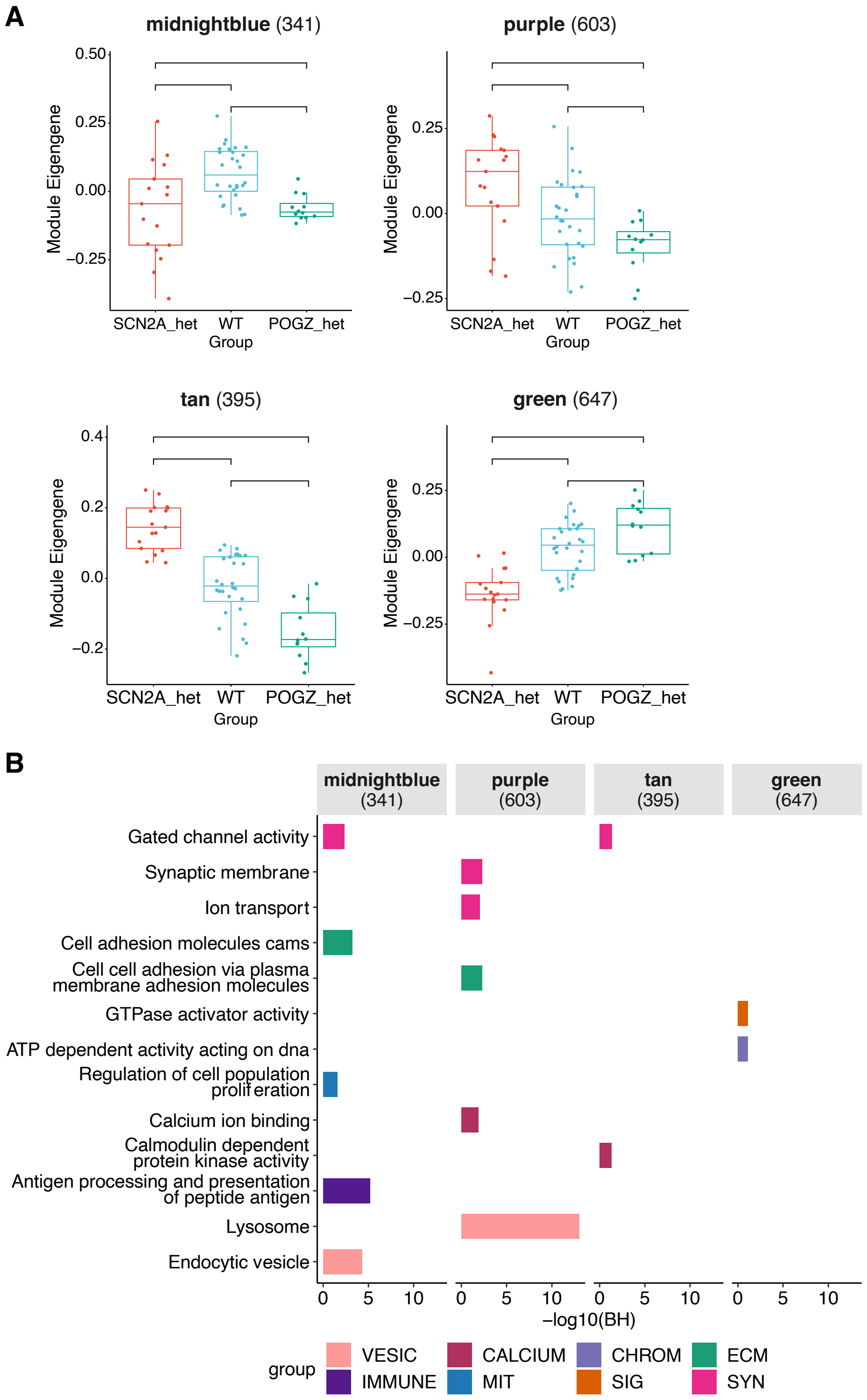


***Supplementary Figure 14. Co-expression modules derived from isogenic heterozygous iN with POGZ or SCN2A gene deletions***

**(A)** Expression levels of eigengenes in gene modules associated with genotype derived from co-expression analysis on isogenic heterozygous iN with *POGZ* or *SCN2A* gene deletions. **(B)** Statistical significance for enrichment of pathways among co-expression modules associated with genotype on heterozygous iN with *POGZ* or *SCN2A* gene deletions. Numbers in parentheses indicate the total number of genes in a given gene list. Colors represent pathway categorization according to the nature of the functional categories related to the enriched term. Groups of functional categories: *VESIC: vesicles, IMMUNE: immune system and inflammation, CALCIUM: calcium signaling, MIT: mitosis and cell proliferation, CHROM: chromatin structure and regulation, SIG: intracellular signaling pathways, ECM: extracellular matrix, SYN: synapse.* BH: Benjamini-Hochberg adjusted p-values.

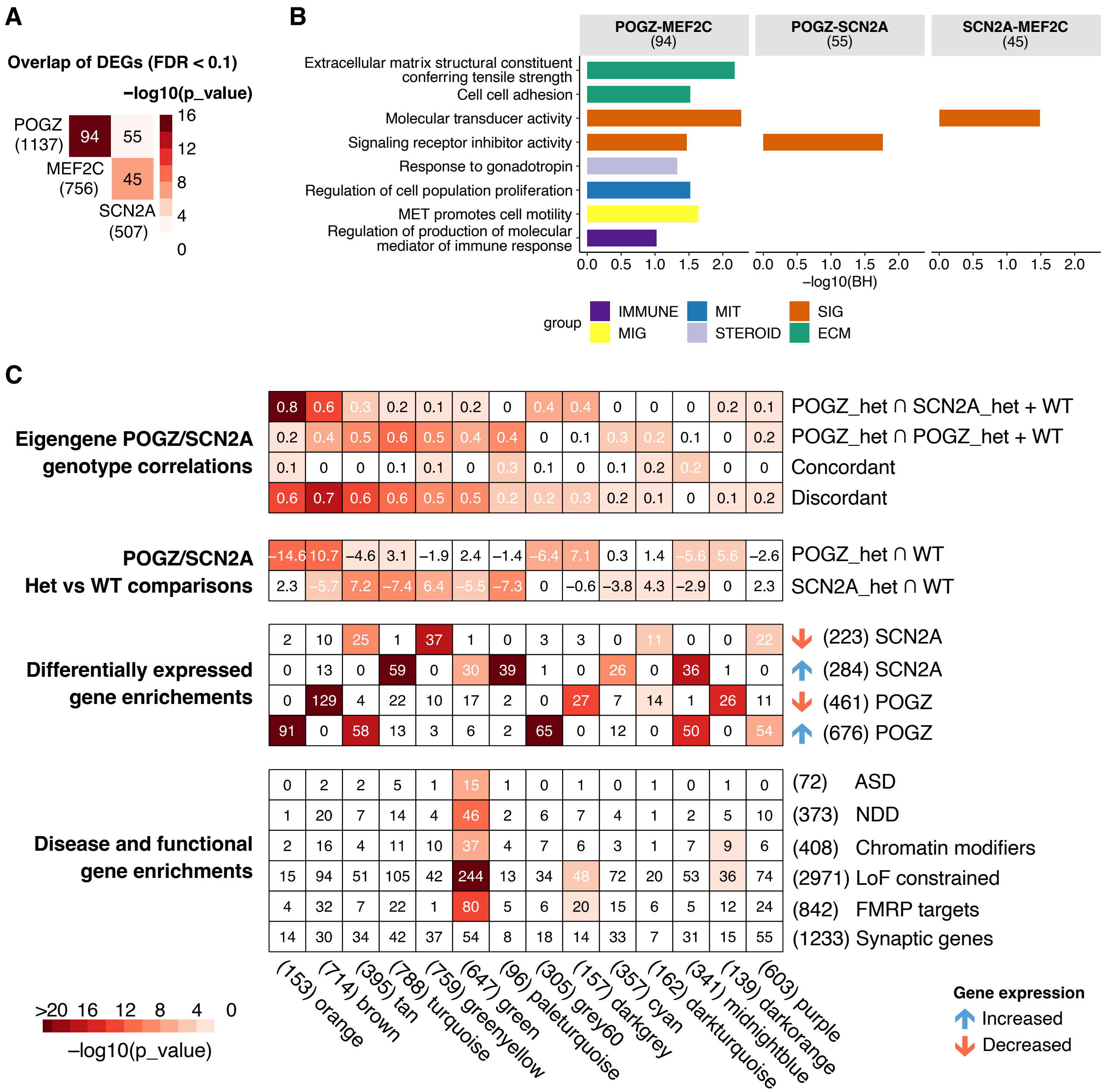


***Supplementary Figure 15. Overlap of co-expression modules derived from isogenic heterozygous iN with POGZ or SCN2A gene deletions***

Heatmap indicating statistical significance for overlap between DEGs (FDR<0.1) from heterozygous iN with *POGZ*, *MEF2C*, or *SCN2A* gene deletions. B. Statistical significance for enrichment of pathways among DEG overlaps from two-way comparisons of heterozygous iN with *POGZ*, *MEF2C* or *SCN2A* gene deletions. Numbers in parentheses indicate the total number of genes in a given gene list. Colors represent pathway categorization according to the nature of the functional categories related to the enriched term. Groups of functional categories: *IMMUNE: immune system and inflammation, MIG: cell migration, MIT: mitosis and cell proliferation, STEROID: steroid metabolism, SIG: intracellular signaling pathways, ECM: extracellular matrix. C. Heatmap indicating statistical significance for genotype model fitting of co-expression analysis from heterozygous iN with POGZ or SCN2A gene deletions. Numbers in parentheses indicate the total number of genes in a given gene list.* BH: Benjamini-Hochberg adjusted p-values.

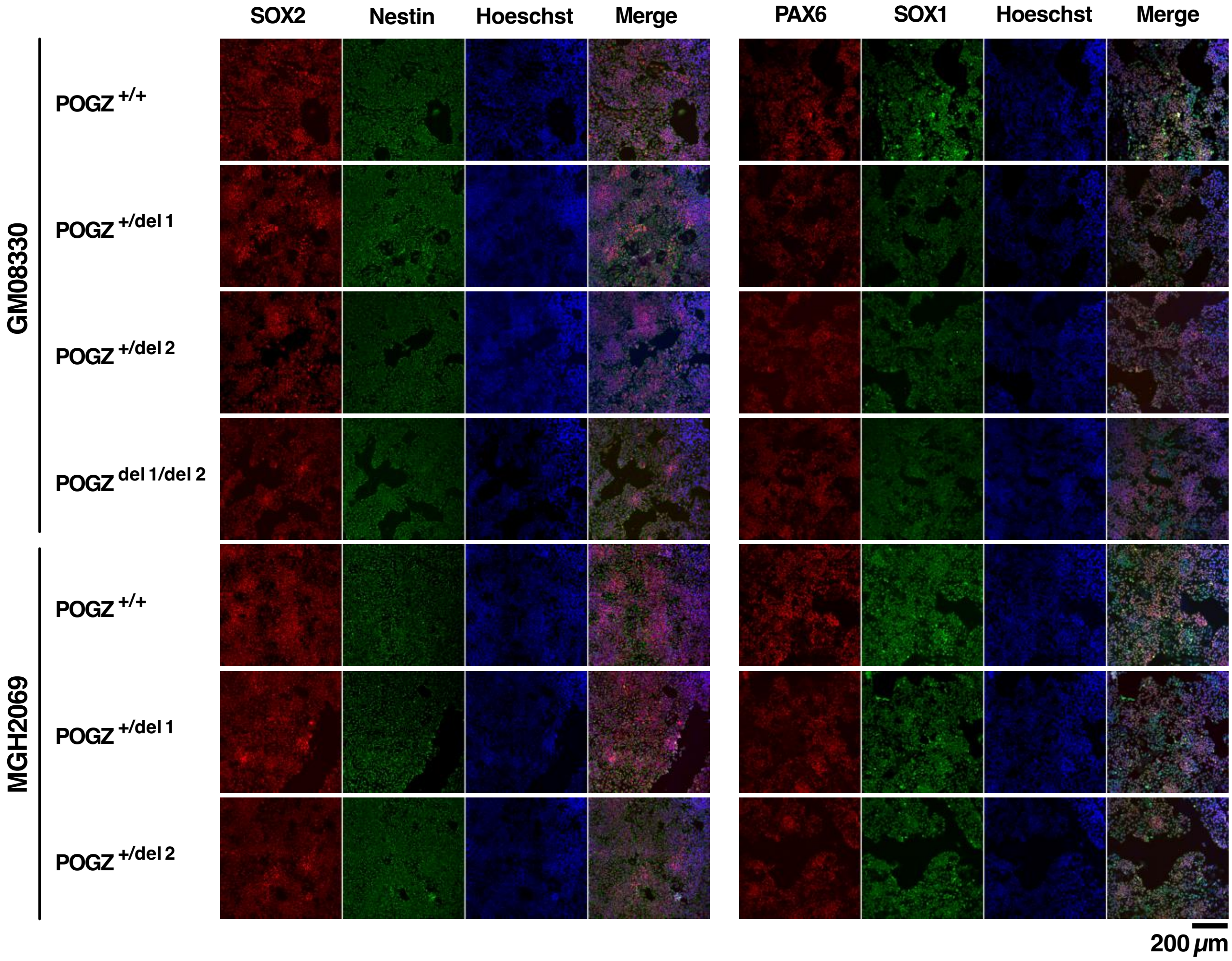


*Supplementary Figure 16. Protein expression validation of neural markers*

At passage 7, WT and edited NSC were plated on coverslips and analyzed for Nestin, PAX6, SOX1 and SOX2 expression by immunostaining using Human Neural Stem Cell Immunocytochemistry Kit (Thermo Fisher Scientific)

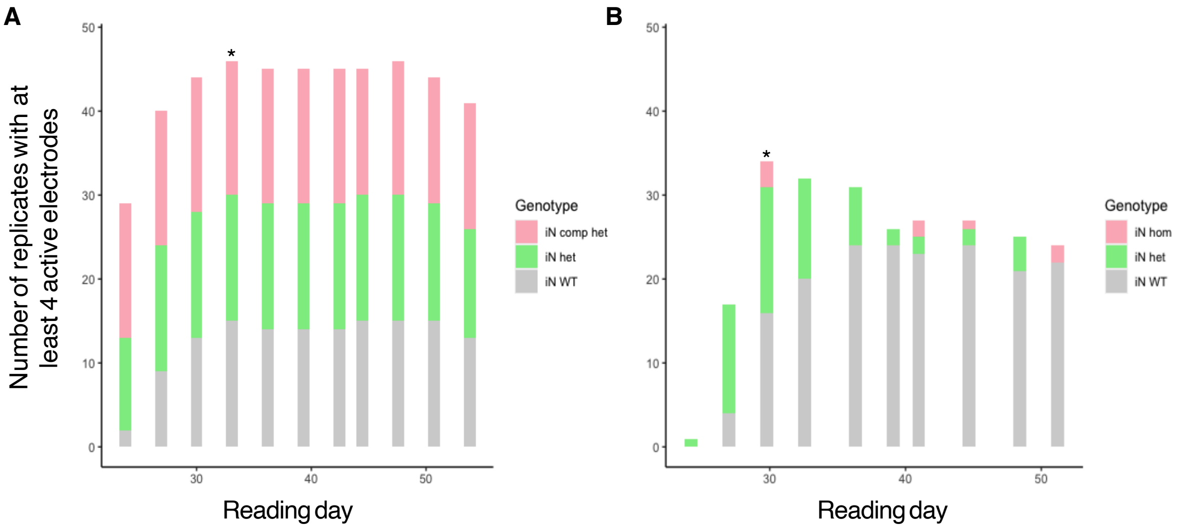


*Supplementary Figure 17. Firing activity over time on iN with POGZ or SCN2A gene deletions.*

**(A)** and **(B)**. Number of iN replicates with at least 4 active electrodes in POGZ (A) or SCN2A (B) Multi-electrode array experiments. Asterisk indicates the reading date with the highest activity rate. *WT: wild type, heterozygous: het, compound heterozygous: comp het, homozygous: hom.*

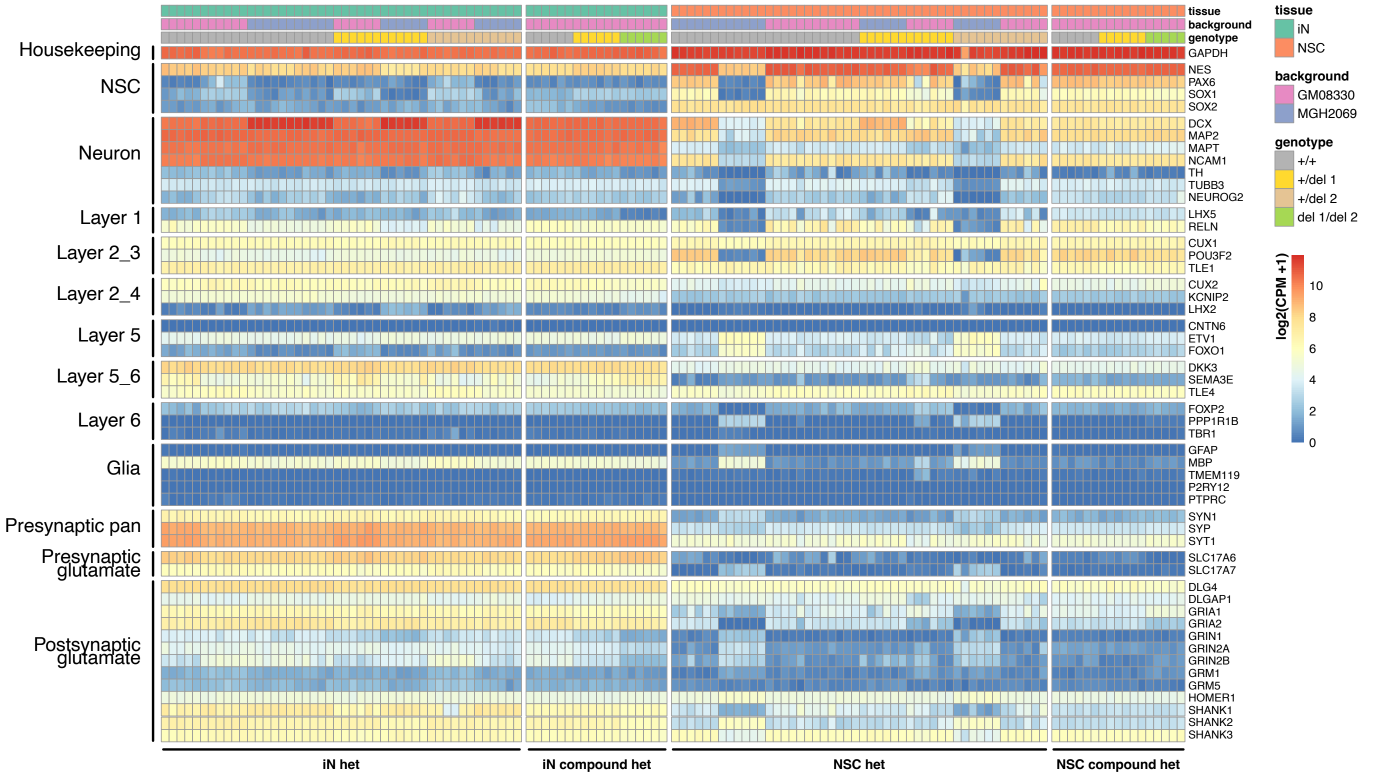


***Supplementary Figure 18. mRNA expression of cell type markers across cell types, background and zygosities***

RNAseq differential analysis considering all annotated *POGZ* transcript isoforms confirmed the expected expression level of the dominant isoforms in heterozygous and compound heterozygous hiPSC-derived neuronal lines (Table S14). To calculate the isoform-specific expression levels of POGZ transcripts, Kallisto [76] was applied to NSC and iN RNAseq libraries, relying on Ensembl GRCh38 v 92 gene and transcript annotations. Kallisto calculated transcriptome-wide expression levels of transcripts in transcript-per-million (tpm). In NSC and iN lines, the top expressed transcript was ENST00000392723, a 6,152 bp transcript encoding a protein of 1,357 amino acids. The same transcript is also the most expressed in human brain tissues and other tissues based on GTEx data[77]. Differential expression analysis based on two-sided t-test revealed that ENST00000392723 was the most significant differentially expressed POGZ transcript, downregulated in heterozygous deletion and compound heterozygous samples in all NSC and iN batches in both backgrounds with p-values varying between 6.6e-3 and 1.75e-6.

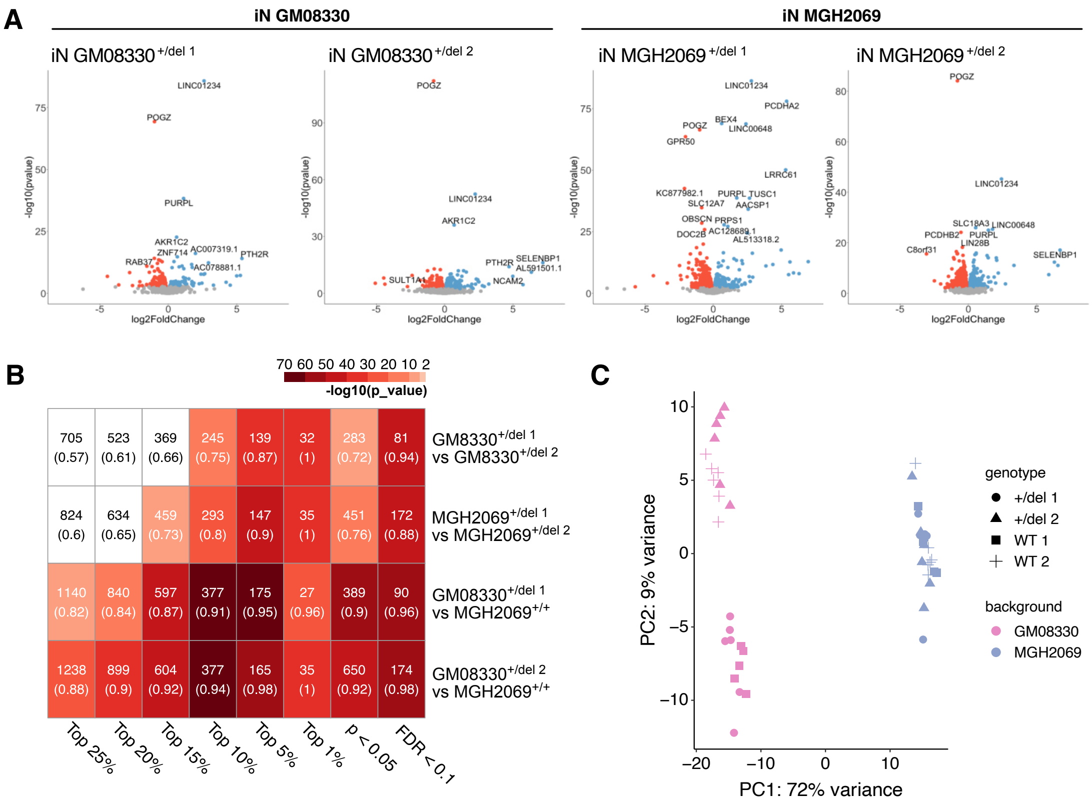


**D**

| Batch | # analyzed | P<0.05 | | | FDR<0.1 | | |
|---|---|---|---|---|---|---|---|
| | | ALL | UP | DOWN | ALL | UP | DOWN |
| $GM08330^{+/del\ 1}$ | 16,769 | 1,486 | 618 | 868 | 269 | 96 | 173 |
| $GM08330^{+/del\ 2}$ | 16,808 | 2,526 | 1,342 | 1,184 | 682 | 389 | 293 |
| $MGH2069^{+/del\ 1}$ | 16,718 | 2,100 | 930 | 1170 | 826 | 311 | 515 |
| $MGH2069^{+/del\ 2}$ | 16,620 | 2,723 | 1,412 | 1,311 | 1,196 | 628 | 598 |

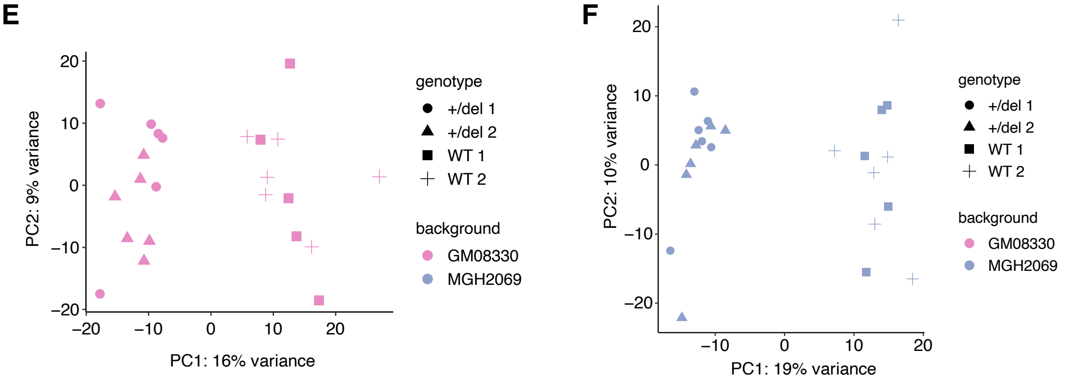


***Supplementary Figure 19 Batch-level differential expression analysis of del1, del2 and their corresponding wild-type iN samples***
**(A)** Volcano plots showing differentially expressed upregulated (blue) and downregulated (red) genes at FDR < 0.1. Not-significant genes (FDR ≥ 0.1) were colored by gray. **(B)** Heatmap highlighting statistical significance of concordant overlap among differentially expressed genes (DEGs) observed between pairs of iN batches. DEGs were identified at different thresholds including FDR < 0.1, p < 0.05 and top 1% to top 25% of p-value ranked lists. Numbers and ratios within parentheses in the cells indicate the number of overlapping DEGs and their ratios to all the overlapping DEGs at a particular threshold for a particular pair of iN batches respectively. Concordantly overlapping DEGs have the same direction of regulation (the same sign of log2FoldChanges). **(C)** Principal component analysis of samples from four iN batches relying on uncorrected expression values. **(D)** Table showing differential expression results for four iN batches. **(E)** and **(F)** Principal component analysis of iN samples from GM08330 background (light pink, E panel) and MGH2069 background (grayish blue, F panel) relying on SVA-corrected expression values.

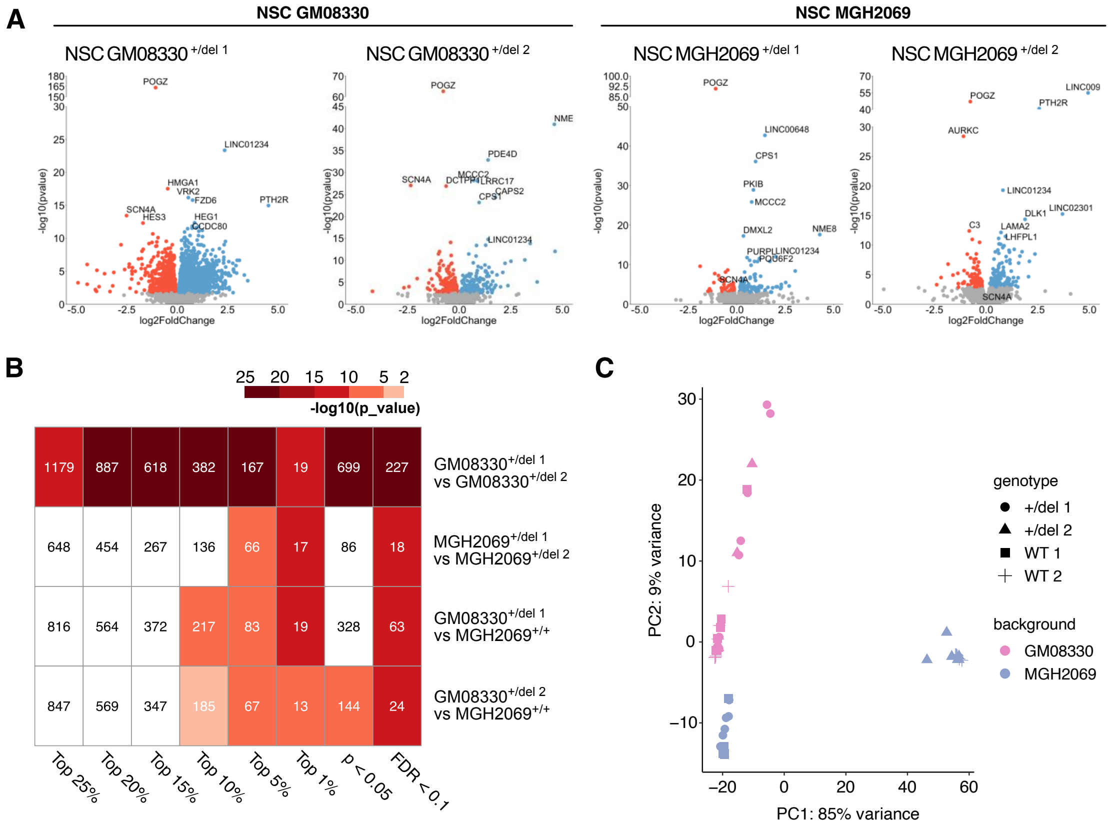


**D**

| Batch | # analyzed | P<0.05 ALL | P<0.05 UP | P<0.05 DOWN | FDR<0.1 ALL | FDR<0.1 UP | FDR<0.1 DOWN |
|---|---|---|---|---|---|---|---|
| GM08330$^{+/del\ 1}$ | 16,681 | 4,440 | 2,653 | 1,787 | 3,277 | 2,022 | 1,255 |
| GM08330$^{+/del\ 2}$ | 16,245 | 1,791 | 885 | 906 | 423 | 204 | 219 |
| MGH2069$^{+/del\ 1}$ | 16,472 | 1,295 | 729 | 566 | 161 | 114 | 47 |
| MGH2069$^{+/del\ 2}$ | 17,543 | 1,179 | 621 | 558 | 238 | 144 | 94 |

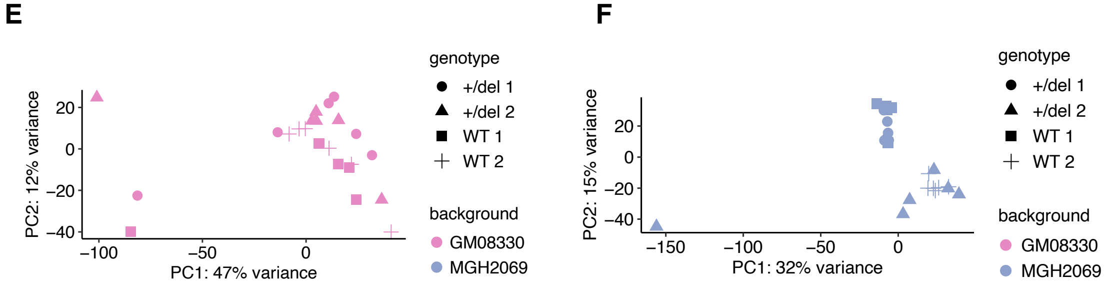


***Supplementary Figure 20. Batch-level differential expression analysis of del1, del2 and their corresponding wild-type NSC samples***
**(A)** Volcano plots showing differentially expressed upregulated (blue) and downregulated (red) genes at FDR < 0.1. Not-significant genes (FDR ≥ 0.1) were colored by gray. **(B)** Heatmap highlighting statistical significance of concordant overlap among differentially expressed genes (DEGs) observed between pairs of NSC batches. DEGs were identified at different thresholds including FDR < 0.1, p < 0.05 and top 1% to top 25% of p-value ranked lists. Numbers and ratios within parentheses in the cells indicate the number of overlapping DEGs and their ratios to all the overlapping DEGs at a particular threshold for a particular pair of NSC batches respectively. Concordantly overlapping DEGs have the same direction of regulation (the same sign of log2FoldChanges). **(C)** Principal component analysis of samples from four NSC batches relying on uncorrected expression values. **(D)** Table showing differential expression results for four NSC batches. **(E)** and **(F)** Principal component analysis of NSC samples from GM08330 background (light pink, E panel) and MGH2069 background (grayish blue, F panel) relying on SVA-corrected expression values.

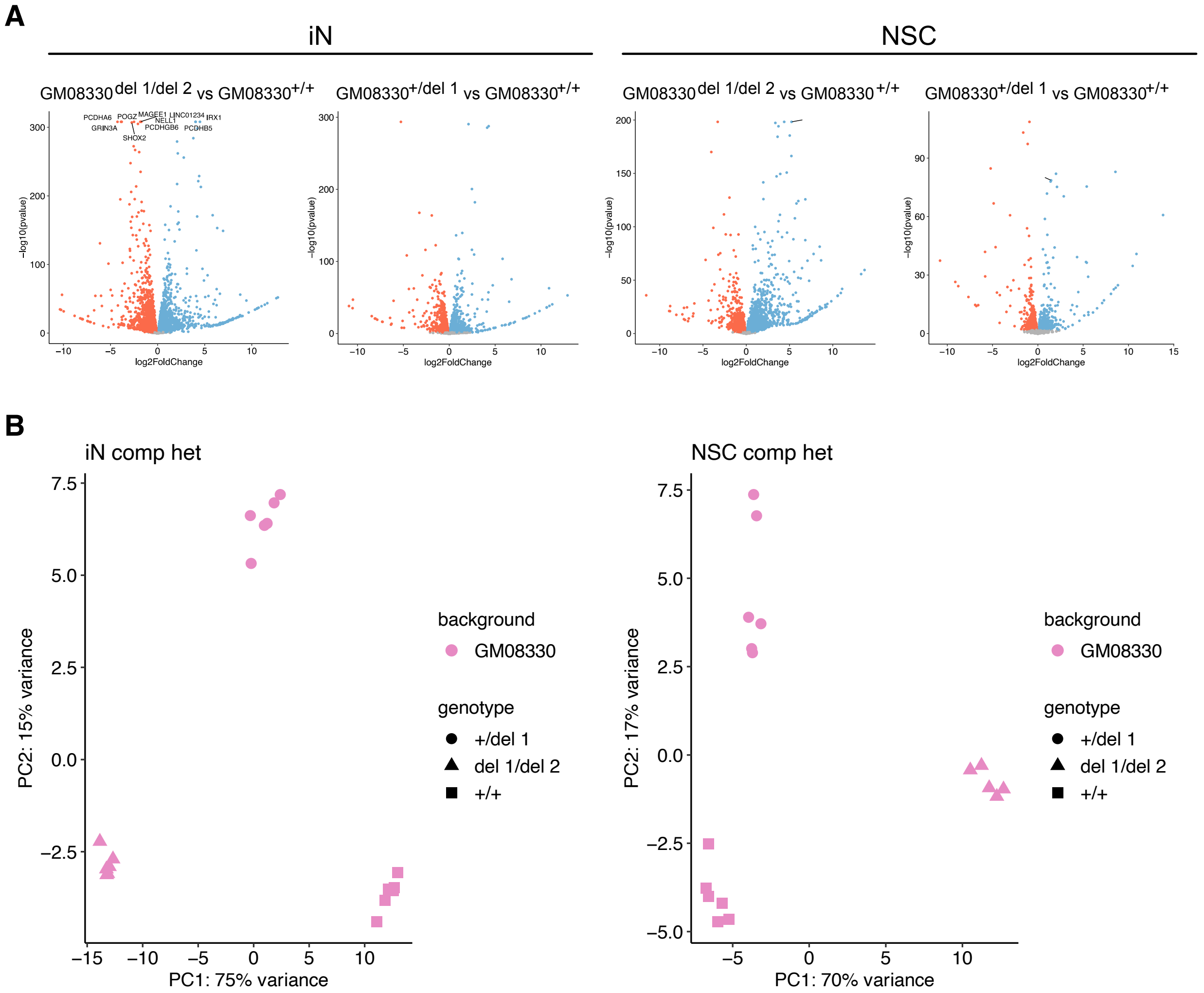


C

| Batch | # analyzed | P<0.05 | | | FDR<0.1 | | |
|---|---|---|---|---|---|---|---|
| | | ALL | UP | DOWN | ALL | UP | DOWN |
| iN GM08330$^{del\ 1/del\ 2}$ | 16,830 | 8,923 | 4,568 | 4,355 | 9,006 | 4,609 | 4,397 |
| iN GM08330$^{+/del\ 1}$ | 16,830 | 4,442 | 2,232 | 2,210 | 3,484 | 1,747 | 1,737 |
| NSC GM08330$^{del\ 1/del\ 2}$ | 16,845 | 5,804 | 3,066 | 2,738 | 5,148 | 2,763 | 2,385 |
| NSC GM08330$^{+/del\ 1}$ | 16,845 | 3,439 | 1,815 | 1,624 | 2,013 | 1,047 | 966 |

***Supplementary Figure 21. Differential expression analysis of compound heterozygous NSC and iN samples***
**(A)** Volcano plots showing differentially expressed upregulated (blue) and downregulated (red) genes at FDR < 0.1. Not-significant genes (FDR ≥ 0.1) were colored by gray. **(B)** Principal component analysis of del1, compound het and wild-type iN samples. **(C)** Principal component analysis of del1, compound het and wild-type NSC samples. **(D)** Table showing differential expression results of compound het vs wild type and del1 vs wild type comparisons in iN and NSC.

A

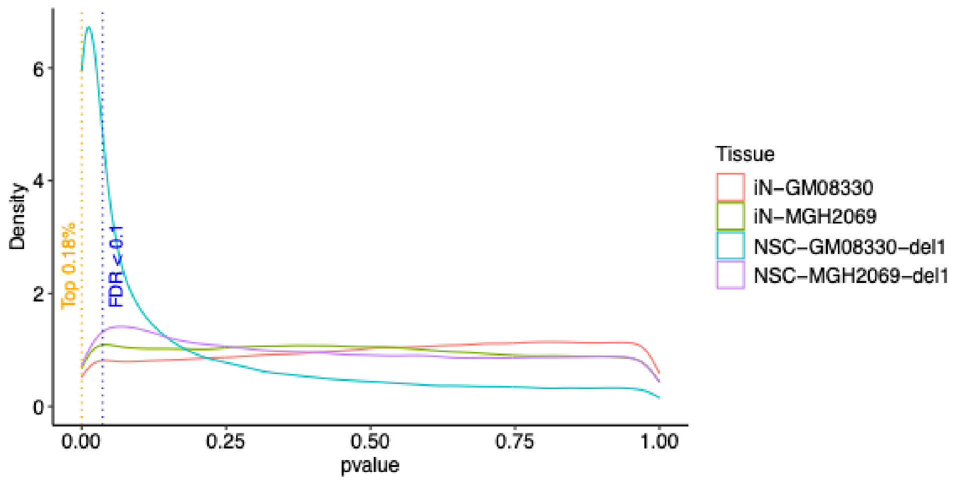


B

**DAR mapped to gene promoters (DAR genes) vs. POGZ targets DEG**

| iN GM80330 (p= 1) | DAR genes | Not DAR genes |
|---|---|---|
| POGZ target DEG | 0 | 160 |
| Non-POGZ target DEG | 13 | 964 |

| iN MGH2369 (p=0.4331) | DAR genes | Not DAR targets |
|---|---|---|
| POGZ target DEG | 4 | 327 |
| Non-POGZ target DEG | 17 | 1738 |

| NSC GM80330 (p=0.7564) | DAR genes | Not DAR genes |
|---|---|---|
| POGZ target DEG | 3 | 1161 |
| Non-POGZ target DEG | 14 | 4119 |

| NSC MGH2369 (p=0.5831) | DAR genes | Not DAR targets |
|---|---|---|
| POGZ target DEGs | 7 | 876 |
| Non-POGZ target DEG | 23 | 2840 |

| iN - backgrounds combined (p= 1) | DAR genes | Not DAR genes |
|---|---|---|
| POGZ target DEG | 0 | 47 |
| Non-POGZ target DEG | 7 | 370 |

| NSC - backgrounds combined (p= 1) | DAR genes | Not DAR targets |
|---|---|---|
| POGZ target DEGs | 0 | 273 |
| Non-POGZ target DEG | 4 | 838 |

***Supplementary Figure 22. Integration between differential expression analysis, chromatin accessibility analysis and POGZ regulatory targets.***
**(A)** Distribution of differentially accessible regions (DAR) according to p-value in differential analysis comparing WT and heterozygous models. hiPSC backgrounds were considered separately. **(B)** Contingency tables for enrichment of differentially accessible regions (DAR) on promoters of differentially expressed POGZ targets. In the columns, genes were categorized as with or without DAR mapped to promoters (“DAR genes” and “Not DAR genes”, respectively). In the rows, differentially expressed genes were categorized as a POGZ target or a non-POGZ target. In each cell type, analysis was performed for each background separately (MGH2069 and GM08330) and combining both backgrounds.

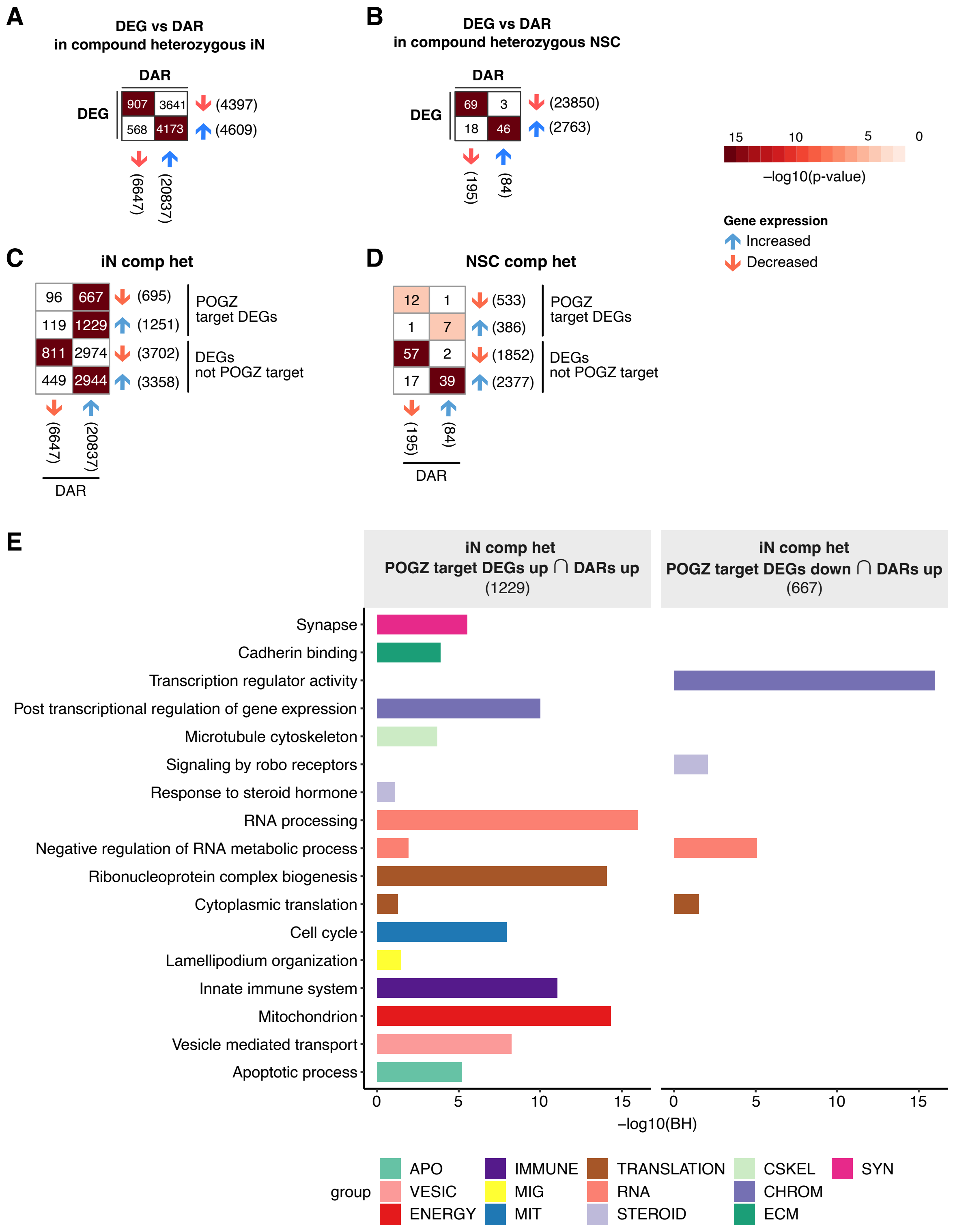


***Supplementary Figure 23. Integration between differential expression and DNA accessibility analysis indicating genes with transcriptional and chromatin accessibility modulation in compound heterozygous lines***

**(A)** and **(B)**. Heatmap indicating statistical significance for overlap between DEGs and DAR-associated genes from human iN (panel A) or NSC (panel B) lines with compound heterozygous gene deletion. **(C)** and **(D)**. Heatmaps indicating statistical significance for overlap of iN (C) or NSC (D) up/ downregulated DEGs and DARs from compound heterozygous lines. E. Statistical significance for enrichment of pathways among POGZ targets that are upregulated (left panel) or downregulated (right panel) DEG and are also under promoters with increased accessibility. Colors represent pathway categorization according to the nature of the biological process related to the enriched term. Numbers in parentheses indicate the total number of genes in a given gene list. *Groups of biological processes: APO: apoptosis, VESIC: vesicles, VASC: vascular system, ENERGY: energy metabolism, IMMUNE: immune system and inflammation, MIG: cell migration, MIT: mitosis and cell proliferation, TRANSLATION: translation, RNA: RNA metabolism, STEROID: steroid metabolism, CSKEL: cytoskeleton, CHROM: chromatin structure and regulation, ECM: extracellular matrix, SYN: synapse.* BH: Benjamini-Hochberg adjusted p-values.

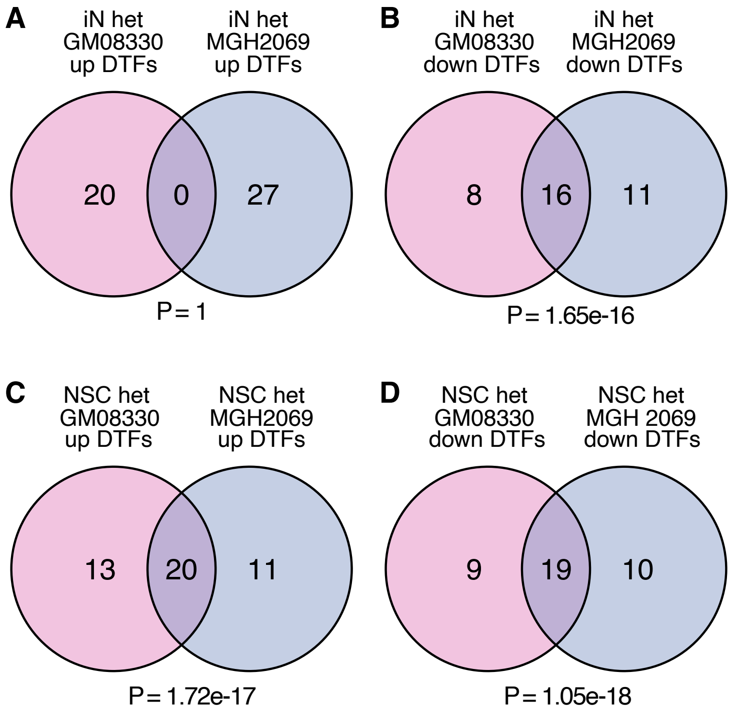


***Supplementary Figure 24. Overlap between DTFs in different background lines and zygosities***

**(A)** to **(D)**. Overlap of DEGs in concordant direction of effect between GM (pink) and MGH (blue) background lines in iN (panel A and B) and NSC (panel C and D).

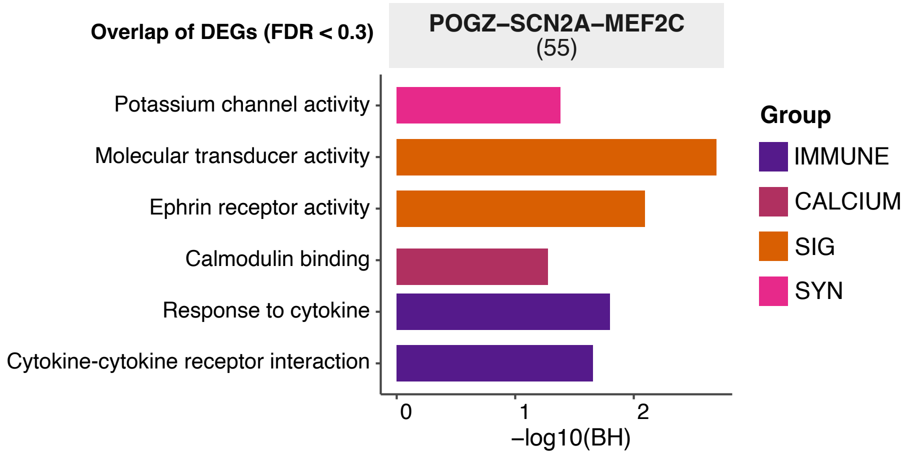


***Supplementary Figure 25. Overlap of transcriptional profiles derived from isogenic heterozygous iN with POGZ, MEF2C or SCN2A gene deletions***

Statistical significance for enrichment of pathways among DEG overlaps from three-way comparisons of heterozygous iN with *POGZ*, *MEF2C* or *SCN2A* gene deletions. Numbers in parentheses indicate the total number of genes in a given gene list. Colors represent pathway categorization according to the nature of the functional categories related to the enriched term. BH: Benjamini-Hochberg adjusted p-values.
